# Comprehensive Chromosome End Remodeling During Programmed DNA Elimination

**DOI:** 10.1101/2020.04.18.047035

**Authors:** Jianbin Wang, Giovana M.B. Veronezi, Yuanyuan Kang, Maxim Zagoskin, Eileen T. O’Toole, Richard E. Davis

## Abstract

Germline and somatic genomes are typically the same in a multicellular organism. However, in some organisms including the parasitic nematode *Ascaris*, programmed DNA elimination leads to a reduced somatic genome compared to germline cells. Previous work on the parasitic nematode *Ascaris* demonstrated that programmed DNA elimination encompasses high fidelity chromosomal breaks at specific genome locations and loss of specific genome sequences including a major tandem repeat of 120 bp and ~1,000 germline-expressed genes. However, the precise chromosomal locations of the 120 bp repeats, the breaks regions, and the eliminated genes remained unknown. Here, we used PacBio long-read sequencing and chromosome conformation capture (Hi-C) to obtain fully assembled chromosomes of *Ascaris* germline and somatic genomes, enabling a complete chromosomal view of DNA elimination. Surprisingly, we found that all 24 germline chromosomes undergo comprehensive chromosome end remodeling with DNA breaks in their subtelomeric regions and loss of distal sequences including the telomeres at both chromosome ends. All new *Ascaris* somatic chromosome ends are recapped by *de novo* telomere healing. We provide an ultrastructural analysis of DNA elimination and show that *Ascaris* eliminated DNA is incorporated into many double membrane-bound structures, similar to micronuclei, during telophase of a DNA elimination mitosis. These micronuclei undergo dynamic changes including loss of active histone marks and localize to the cytoplasm following daughter nuclei formation and cytokinesis where they form autophagosomes. Comparative analysis of nematode chromosomes suggests that germline chromosome fusions occurred forming *Ascaris* sex chromosomes that become independent chromosomes following DNA elimination breaks in somatic cells. These studies provide the first chromosomal view and define novel features and functions of metazoan programmed DNA elimination.

## Introduction

Genomes in the diverse cells of a metazoan are typically the same. A variety of mechanisms have evolved to maintain and ensure this genome constancy. Changes in the genome, particularly rearrangements or significant sequence loss, can be deleterious leading to disease or inviability. However, there are exceptions to this genome constancy, including gene rearrangements that occur in B-cells and T-cells that expand the repertoire of immunoglobulins and receptors, respectively [1]. Another major exception is programmed DNA elimination. It occurs in a breadth of evolutionarily diverse organisms ranging from ciliates to mammals [2]. Metazoan DNA elimination occurs primarily in two major forms. In the first form, chromosome loss, one or many chromosomes are eliminated. This form occurs in a few insects, mites, nematodes (*Strongyloides ratti*), hagfish, frogs, songbirds and marsupials [2–5], and it is typically associated with sex determination or species that reproduce by hybridogenesis [2,6,7]. In the second form, chromosomes break with retention of some chromosome fragments and the elimination of others. The elimination only occurs in somatic cells and leads to a distinct and reduced somatic cell genome compared to the germline cell genome. This form of DNA elimination occurs in some parasitic nematodes (ascarids), copepods, ratfish, hagfish, and lampreys [4,8–16]. Programmed DNA rearrangements also occur in ciliates distinguishing the germline from somatic nuclei [17–21]. However, aspects of DNA elimination in metazoa appears mechanistically distinct from that described in ciliates.

Programmed DNA elimination in ascarid parasitic nematodes results in the loss of 13-90% of the germline DNA in forming the somatic genome [16,22]. The majority of eliminated DNA are repetitive sequences organized in tandem repeats that differ in the three genera examined (*Ascaris suum* = 120 bp repeat, *Parascaris univalens* = 5 and 10 bp repeats, and *Toxocara canis* = 49 bp repeat). One thousand (*Ascaris* and *Parascaris*) to two thousand genes (*Toxocara*) are also eliminated representing 5-10% of all genes, respectively. These genes are primarily expressed in the germline and early embryos. Comparison of the conservation of eliminated genes among these three nematodes indicates that the genes are expressed predominantly during spermatogenesis. We have proposed that DNA elimination in nematodes is a mechanism for silencing germline expressed genes [16,22]. Instead of using epigenetic mechanisms to silence germline genes in somatic cells, these genes are silenced by their elimination from the genome.

Ascarid programmed DNA elimination involves chromosome breaks, *de novo* telomere healing of the breaks, and the selective retention and loss of chromosome fragments. Previous analysis of the DNA breaks, called chromosomal break regions (CBRs) [23–25], demonstrated they occur with high fidelity in 5 independent pre-somatic lineages within regions of 3-6 kb [16,22]. While the CBRs are high fidelity at the chromosome level (3-6 kb within 10 Mb chromosomes), the telomere healing occurs randomly within the CBRs. It is currently not known whether this heterogeneity within the break regions is due to heterogeneity in DNA breaks or telomere healing. Extensive analyses of CBRs did not identify any sequence, structural motifs, histone marks, small RNAs, or other features that define or target these regions as sites for the DNA breaks in chromosomes [16]. CBRs, however, become more open or chromatin accessible based on ATAC-seq just prior to and during DNA elimination [16]. *Ascaris* has holocentric chromosomes with centromeres/kinetochores extending along the length of the chromosomes in gametogenic germline cells [26–28]. Prior to DNA elimination, regions of chromosomes that will be lost undergo a reduction of CENPA and kinetochores [28]. During a DNA elimination mitosis, chromosome fragments with reduced CENPA and kinetochores remain at or near the metaphase plate during anaphase, are not segregated into the newly formed daughter nuclei, and become localized to the cytoplasm following telophase and cytokinesis where the DNA is eventually degraded. Thus, both discrete sites for DNA breaks and chromosomal changes in CENP-A (centromeres/kinetochores) localization are involved in DNA elimination and define which chromosome fragments will be retained or eliminated.

Our previous work on *Ascaris* DNA elimination identified 40 CBRs and the eliminated DNA [16]. However, the previous chromosome assemblies were not sufficient to define the ends of the chromosomes or a comprehensive view of the chromosomal location of the DNA breaks (CBRs) and eliminated DNA. Here, we used in-depth PacBio Sequel II long read sequencing and chromosome conformation capture (Hi-C) to generate comprehensive chromosomal assemblies of *Ascaris* germline and somatic chromosomes. The new genome assemblies enabled us to define the subtelomeric/telomeric regions of all the 24 germline chromosomes and identify the chromosomal location of all CBRs (72 total with 32 new CBRs identified) in the 24 chromosomes, the retained and eliminated DNA regions, and the location and organization of repetitive elements and eliminated genes. DNA FISH and comparative genome analysis of the ends of germline and somatic chromosome assemblies demonstrated that during DNA elimination both ends of all 24 germline chromosomes undergo subtelomeric DNA breaks (48 total DNA breaks), *de novo* telomere healing of the chromosome ends, and the loss of distal chromosome sequences. These data indicate that all *Ascaris* chromosome ends undergo significant remodeling during DNA elimination. DNA breaks at 24 internal sites generates additional smaller chromosomes with an increase in chromosome number from 24 in the germline to 36 in the somatic cells. Electron microscopy studies reveal an ultrastructural timeline of DNA elimination where diffuse chromatin is first identified in the spindle proper and the metaphase plate in late prometaphase; this chromatin is less condensed than the other chromosomes and does not assemble a kinetochore/KMTs. As such, the diffuse chromatin is not segregated during anaphase and remains at the spindle midplane. During telophase of DNA elimination mitoses, the eliminated DNA is incorporated and segregated into double membrane-bound structures that are reminiscent of micronuclei. Eliminated DNA once incorporated into micronuclei undergoes chromatin changes and form autophagosomes in the cytoplasm. Our data also reveal that recent germline chromosome fusion events contributed to the evolution of *Ascaris* sex chromosomes; CBRs and DNA elimination separate these fusions into independent somatic chromosomes. These studies provide the most comprehensive genome analysis of DNA elimination in a metazoan, an ultrastructural analysis of DNA elimination, and identifies additional novel and key aspects of metazoan DNA elimination.

## Results

### PacBio and Hi-C enable complete genome assembly

Our previous studies established a reference genome for the *Ascaris* germline [16]. However, due to the large 120 bp tandem repeat clusters and other complex repeats including the subtelomeric regions, the technologies used (Illumina, PacBio [average 10 kb reads], and optical mapping) were not sufficient to assemble a comprehensive full-chromosome genome. To overcome this, we generated ~360X raw PacBio Sequel II long reads with ~100X read coverage over 40 kb in length (average 52 kb). The PacBio reads were assembled into 763 contigs (N50 = 4.83 Mb) using Canu [29]. To further scaffold these contigs into chromosomes, we obtained Hi-C data from both germline and somatic cells. The Hi-C data enabled us to assemble the germline genome into 24 germline chromosomes (N50 = 12.43 Mb) with only 2.3% of the genome in 84 unplaced contigs (see Supplementary Material for a detailed description of the genome assembly). Our somatic Hi-C data and genome assembly indicates that the germline genome is reduced and split into 36 somatic chromosomes with no apparent genome recombination or rearrangement (Fig. 1).

**Figure 1.**
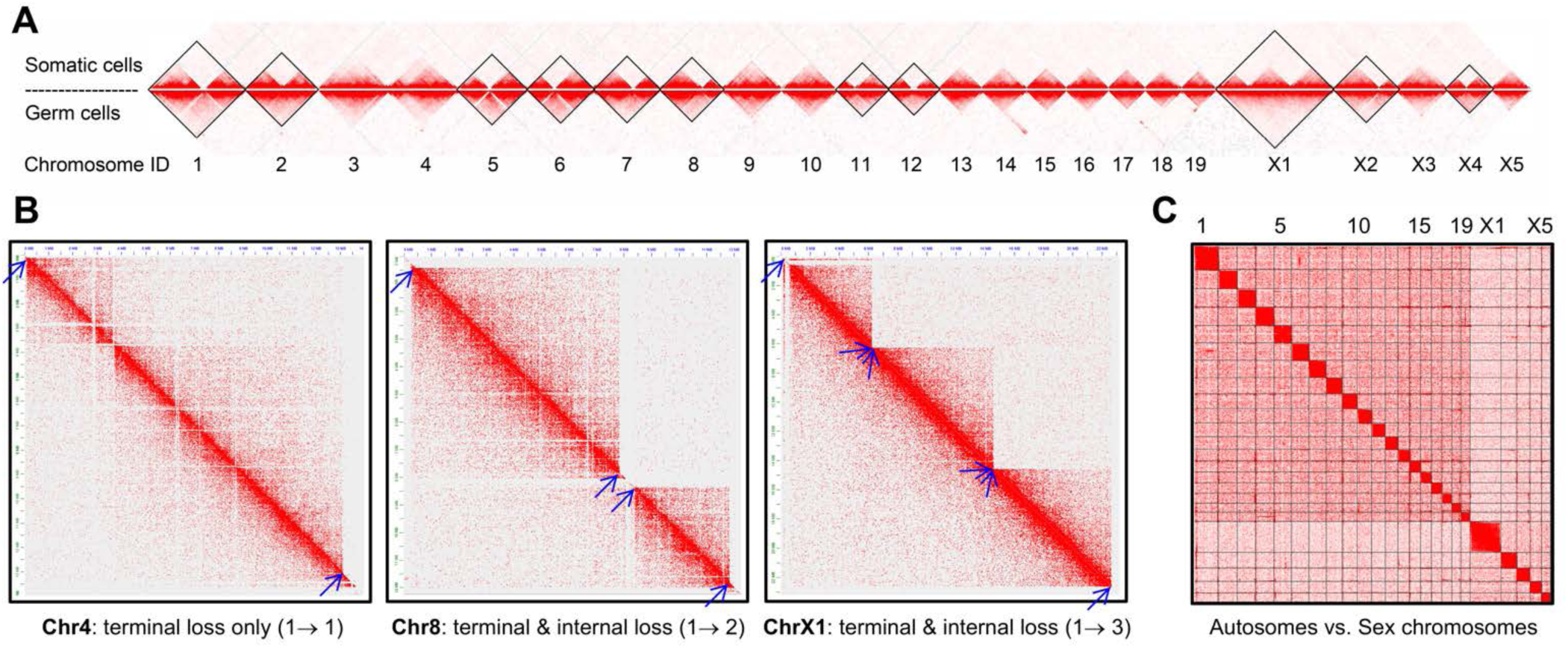
*Ascaris* chromosome Hi-C interactions. **A** and **B**. Hi-C interactions in germline and somatic cell genomes (**A:** overview of all chromosomes; **B:** selected examples), illustrating the chromosomal changes before and after *Ascaris* DNA elimination. In **A**, Hi-C data from germline (testis, bottom) and somatic cells (5 day or 32-64 cell embryos, top) were mapped to the 24 assembled *Ascaris* germline chromosomes. The density of the heatmap (Juicebox v1.11.08) indicates the relative contact probability of two loci. The strongest interactions are within chromosomes as illustrated in the 24 germline (bottom) or 36 somatic blocks that correspond to chromosomes (top). Eleven chromosomes (marked in black rhombus boxes) undergo breaks during DNA elimination leading to an increase in chromosomes. No chromosomal rearrangements were identified in the somatic Hi-C data. In **B**, selected germline chromosomes with the Hi-C interactions are shown (bottom left = germline and top right = somatic). These examples are for germline chromosomes with no internal chromosome changes (Chr4) or increased chromosomes following DNA elimination (Chr8 and ChrX1). Blue arrows point to the CBRs. Note the loss of DNA in the somatic cells at the chromosome ends. See **Figure S1** for details of the Hi-C comparisons in all 24 chromosomes. **C**. Hi-C data showing the interactions between *Ascaris* autosomes and sex chromosomes (X1-X5) in the testis. The dataset from **A** was used to illustrate the interchromosomal interactions among *Ascaris* chromosomes.

We identified 19 autosomes and 5 sex chromosomes based on the read coverage for the 24 germline chromosomes (see Fig. 1 and Table 1). The 5 sex chromosomes are consistent with previous cytogenetic analyses [30]. Hi-C data indicate the sex chromosomes in the testis have greater interactions with each other than with the autosomes (Fig. 1C). This high level of interaction among the sex chromosomes suggests that they are likely physically close to each other during meiosis, reminiscent of the chromosome behavior during meiosis in other systems with multiple sex chromosomes (e.g., tiger beetles and platypus) [31–33]. The level of interaction among these sex chromosomes is reduced between autosomes and sex chromosomes in 32-64 cell embryo somatic cells (Supplementary Figure S2), suggesting these sex chromosomes are more intermingled with autosomes in the somatic cells.

We identified 34 genomic loci in 22 chromosomes for the major 120 bp repeat clusters (hereafter called 120 bp clusters) (Table 1 and Fig. 2). Most of these clusters exceed the length of the PacBio long reads (50-100 Kb). As the total amount of the eliminated 120 bp DNA is around 30 Mb, we estimate that on average each 120 bp cluster is ~900 Kb. PacBio reads of these clusters consist primarily of 120 bp repeats and its variants. However, other repetitive sequences can be interspersed and/or flank these 120 bp repeats. Most of the 120 bp clusters are in subtelomeric regions (25) of the chromosomes, while 9 are in internal regions of the chromosomes (Table 1 and Fig. 2). All of the 120 bp repeat is eliminated from somatic cells; no sequence over 30 bp matching the 120 bp sequence is present in the somatic genome. We were able to fully assemble 6 (out of 48) subtelomeric-telomeric regions for *Ascaris* germline chromosomes. For the remaining subtelomeric-telomeric chromosome ends, analysis of the PacBio reads suggests that at least 50-100 kb of subtelomeric repetitive sequence precedes the telomeres (Fig. 2) precluding their complete assembly. Germline telomere length is on average > 15 kb with many PacBio reads for the telomeres exceeding 50 kb.

**Figure 2.**
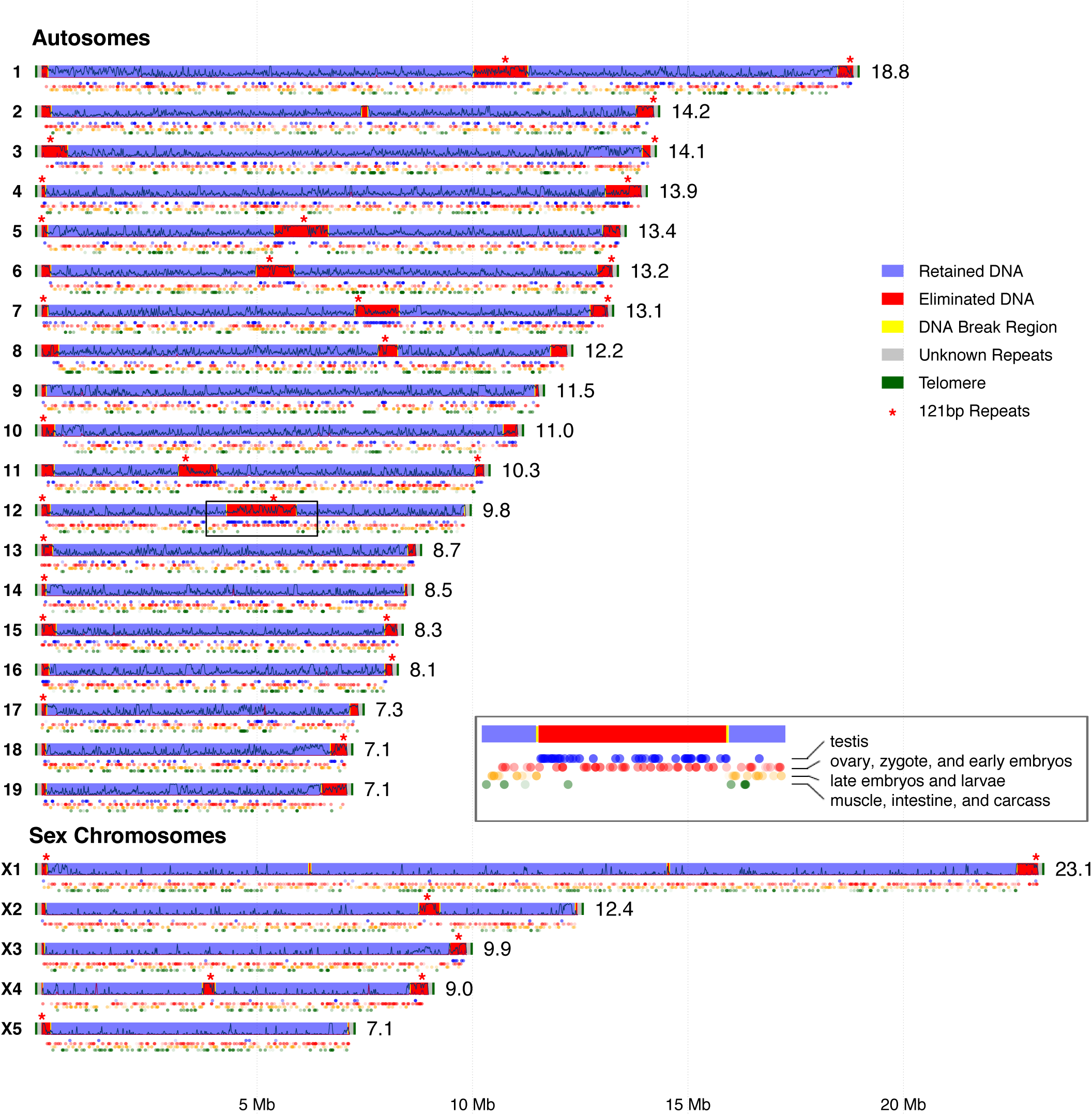
*Ascaris* chromosomes and programmed DNA elimination. Chromosomes are illustrated with the length of the chromosome in Mb on the right. Each dot beneath the chromosomes is a gene with the color and transparency indicating the extent of tissue specific expression (see the boxed enlarged region in chr12 for legend). Over 70% of the genes (with average rpkm >= 2 and maximum rpkm >= 5 across the 9 indicated tissues) were plotted. Tissue specific expression was measured by comparing the gene expression level (rpkm) across the tissues, with the most solid color indicating a rpkm over 4-fold higher than any other tissues (see Methods for details). The amount of repetitive sequences for *Ascaris* chromosomes is illustrated within the chromosome bars, with simple satellite repeats in red and all other repeats in black across the chromosome.

Overall, we have used long PacBio reads and Hi-C to assemble 24 complete *Ascaris* germline chromosomes including the 5 sex chromosomes. Although this genome assembly is not gap free, we have essentially anchored all sequences into the 24 chromosomes, identified the locations for the 34 large 120 bp repeat clusters, and characterized the organization of the 120 bp repeats as well as the subtelomeric repeats. This complete genome assembly provides a significant new resource for an important parasite infecting upwards of 800 million people [34] and enables a chromosomal view of programmed DNA elimination.

**Table 1.**
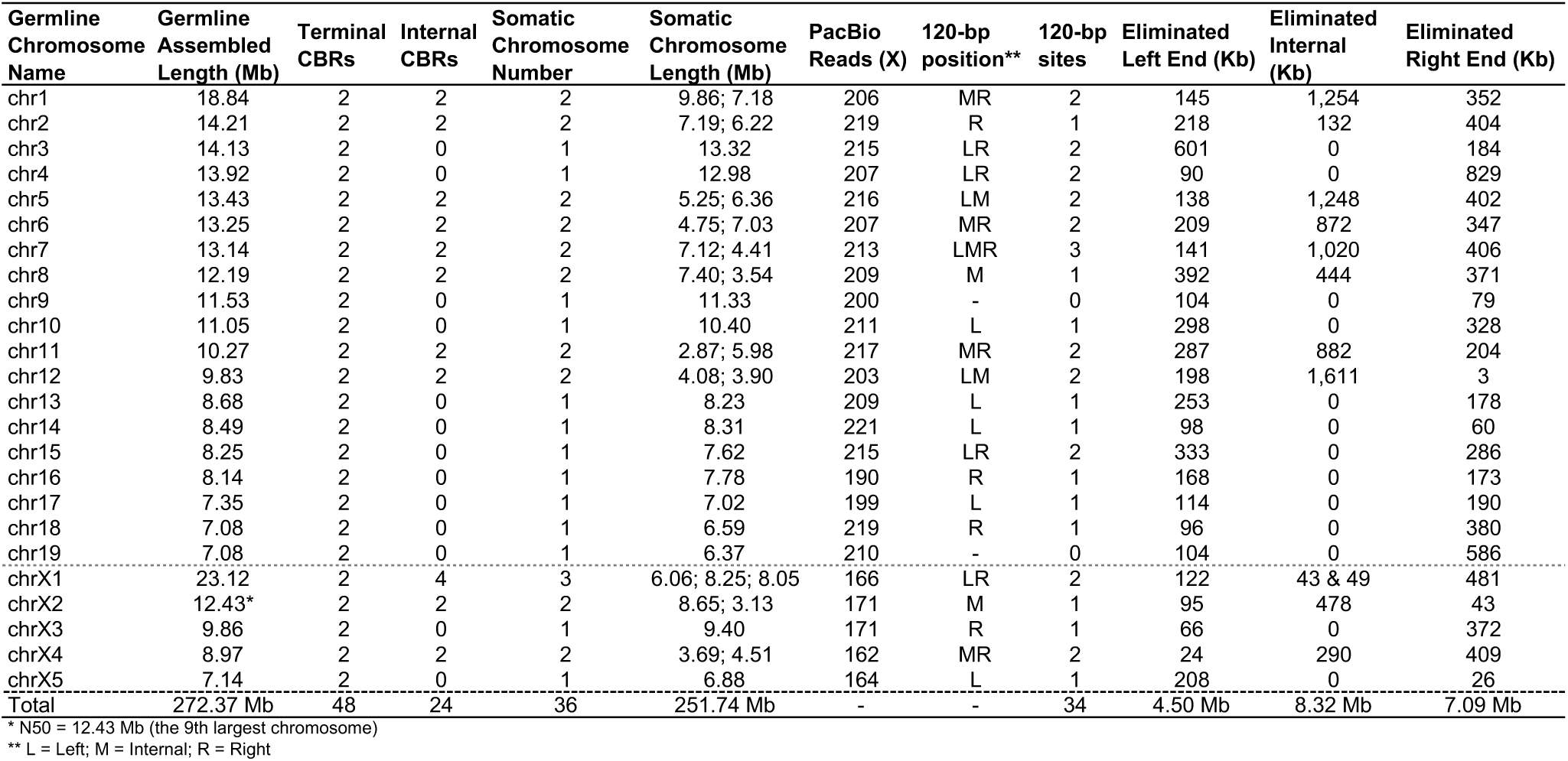
*Ascaris* germline and somatic chromosomes.

| Germline Chromosome Name | Germline Assembled Length (Mb) | Terminal CBRs | Internal CBRs | Somatic Chromosome Number | Somatic Chromosome Length (Mb) | PacBio Reads (X) | 120-bp position** | 120-bp sites | Eliminated Left End (Kb) | Eliminated Internal (Kb) | Eliminated Right End (Kb) |
| --- | --- | --- | --- | --- | --- | --- | --- | --- | --- | --- | --- |
| chr1 | 18.84 | 2 | 2 | 2 | 9.86; 7.18 | 206 | MR | 2 | 145 | 1,254 | 352 |
| chr2 | 14.21 | 2 | 2 | 2 | 7.19; 6.22 | 219 | R | 1 | 218 | 132 | 404 |
| chr3 | 14.13 | 2 | 0 | 1 | 13.32 | 215 | LR | 2 | 601 | 0 | 184 |
| chr4 | 13.92 | 2 | 0 | 1 | 12.98 | 207 | LR | 2 | 90 | 0 | 829 |
| chr5 | 13.43 | 2 | 2 | 2 | 5.25; 6.36 | 216 | LM | 2 | 138 | 1,248 | 402 |
| chr6 | 13.25 | 2 | 2 | 2 | 4.75; 7.03 | 207 | MR | 2 | 209 | 872 | 347 |
| chr7 | 13.14 | 2 | 2 | 2 | 7.12; 4.41 | 213 | LMR | 3 | 141 | 1,020 | 406 |
| chr8 | 12.19 | 2 | 2 | 2 | 7.40; 3.54 | 209 | M | 1 | 392 | 444 | 371 |
| chr9 | 11.53 | 2 | 0 | 1 | 11.33 | 200 | - | 0 | 104 | 0 | 79 |
| chr10 | 11.05 | 2 | 0 | 1 | 10.40 | 211 | L | 1 | 298 | 0 | 328 |
| chr11 | 10.27 | 2 | 2 | 2 | 2.87; 5.98 | 217 | MR | 2 | 287 | 882 | 204 |
| chr12 | 9.83 | 2 | 2 | 2 | 4.08; 3.90 | 203 | LM | 2 | 198 | 1,611 | 3 |
| chr13 | 8.68 | 2 | 0 | 1 | 8.23 | 209 | L | 1 | 253 | 0 | 178 |
| chr14 | 8.49 | 2 | 0 | 1 | 8.31 | 221 | L | 1 | 98 | 0 | 60 |
| chr15 | 8.25 | 2 | 0 | 1 | 7.62 | 215 | LR | 2 | 333 | 0 | 286 |
| chr16 | 8.14 | 2 | 0 | 1 | 7.78 | 190 | R | 1 | 168 | 0 | 173 |
| chr17 | 7.35 | 2 | 0 | 1 | 7.02 | 199 | L | 1 | 114 | 0 | 190 |
| chr18 | 7.08 | 2 | 0 | 1 | 6.59 | 219 | R | 1 | 96 | 0 | 380 |
| chr19 | 7.08 | 2 | 0 | 1 | 6.37 | 210 | - | 0 | 104 | 0 | 586 |
| chrX1 | 23.12 | 2 | 4 | 3 | 6.06; 8.25; 8.05 | 166 | LR | 2 | 122 | 43 & 49 | 481 |
| chrX2 | 12.43* | 2 | 2 | 2 | 8.65; 3.13 | 171 | M | 1 | 95 | 478 | 43 |
| chrX3 | 9.86 | 2 | 0 | 1 | 9.40 | 171 | R | 1 | 66 | 0 | 372 |
| chrX4 | 8.97 | 2 | 2 | 2 | 3.69; 4.51 | 162 | MR | 2 | 24 | 290 | 409 |
| chrX5 | 7.14 | 2 | 0 | 1 | 6.88 | 164 | L | 1 | 208 | 0 | 26 |
| Total | 272.37 Mb | 48 | 24 | 36 | 251.74 Mb | - | - | 34 | 4.50 Mb | 8.32 Mb | 7.09 Mb |

\* N50 = 12.43 Mb (the 9th largest chromosome)
\*\* L = Left; M = Internal; R = Right

### Gene, repeat, and centromere distribution in *Ascaris* chromosomes

The general organization of a nematode autosome based on studies from *C. elegans* [35] indicates that the middle third of the chromosome is enriched for conserved genes and contains fewer repetitive sequences compared to the chromosome “arms” which have fewer numbers of genes and are repeat rich. The genes in the *C. elegans* chromosome “arms” are in general are not well-conserved. In contrast, *Ascaris* autosomes exhibit a relatively uniform distribution of genes with a few gene-poor regions adjacent to subtelomeric regions and the 120 bp repeat clusters, many of which are in internal regions of chromosomes (Fig. 2). Repetitive sequences, with the exception of the 120 bp repeat clusters, are relatively uniformly distributed along the chromosomes except for increased repeat levels in subtelomeric regions. Heterochromatic regions (measured by H3K9me2/3 ChIP-seq; data not shown) are in general associated with the repetitive sequences in *Ascaris* chromosomes. *Ascaris* has holocentromeres and the new assembly indicates that the holocentromere/CENP-A is in general distributed along the full-length of the chromosomes, consistent with our previous findings [27].

*Ascaris* and *C. elegans* have a similar number of genes (18,000 vs. 20,000). However, the *Ascaris* genome is 3 times larger, mainly due to the additional introns and the size of introns. To evaluate the tissue-specific expression of *Ascaris* genes and their chromosomal location, we analyzed our comprehensive RNA-seq data [21,22,36]. Some tissue-specific genes are clustered in the genome, particularly testis-specific genes that are eliminated (Fig. 2, see boxed region on chromosome 12). Genes expressed in the ovary, zygotes, and early embryos are also clustered (for example, chromosome 4 middle, right and chromosome 12 left end) as are some other genes expressed in the soma (chromosome 9 middle, right). Genes on the sex chromosomes are primarily genes expressed in the ovary, zygotes, and early embryos with very few testis specific genes, with the exception of one cluster in the subtelomeric region on sex chromosome X3 that is lost in DNA elimination.

### Chromosomal break regions (CBRs) for DNA elimination

We compared germline and somatic genome coverage to identify regions of the genome that were lost following DNA elimination to form the somatic genome [16,22]. A hallmark of nematode DNA elimination is a chromosome break, the loss of DNA sequences, and the formation of a new chromosome end with a *de novo* telomere in the somatic genome [16,22–25]. We identified 72 sites for DNA breaks (32 new sites) and formation of new telomeres and their chromosomal location. All sites of new telomere addition occur heterogeneously within 3-6 kb regions, known as chromosomal break regions (CBRs) [16,22–25]. The CBRs reproducibly occur with high fidelity at specific locations in the chromosomes in all five pre-somatic cells and individuals that undergo DNA elimination. Our sequence analyses of the new CBRs did not identify any specific sequence or structural features (including motifs, hairpins, palindromes, G-quadruplexes, Z-DNA, or repetitive sequences) within or near the CBRs or small RNAs targeting the regions that might specify or recruit machinery to the CBRs as previously described [16,22–25]. However, as for the previously described CBRs, the newly identified CBRs all become more chromatin accessible (measured by ATAC-seq) just before the DNA elimination mitosis [16] (see Supplementary Figure S3).

### DNA elimination remodels the ends of all germline chromosomes

DNA FISH using a telomere probe indicated that telomeres on all chromosomes are reduced below detection during a DNA elimination mitosis (Figure 3A-C). Telomere re-extension seems to be heterogeneous and take multiple cell cycles to reach signal intensity similar to the germline cells (Figure 3D-E). Similar observations were made previously by Niedermaier and Moritz [30]. Our initial interpretation of these data was that the telomeres may have undergone a cataclysmic recombination event that removed most of the telomere while leaving a small portion of the telomere on the chromosome that was below the level of detection of DNA FISH hybridization [37–39]. However, our experimental efforts using a variety of approaches (Southern Blots, STELA, PacBio sequencing, data not shown) [40] were not able to confirm that the telomeres were shortened on the ends of chromosomes.

**Figure 3.**
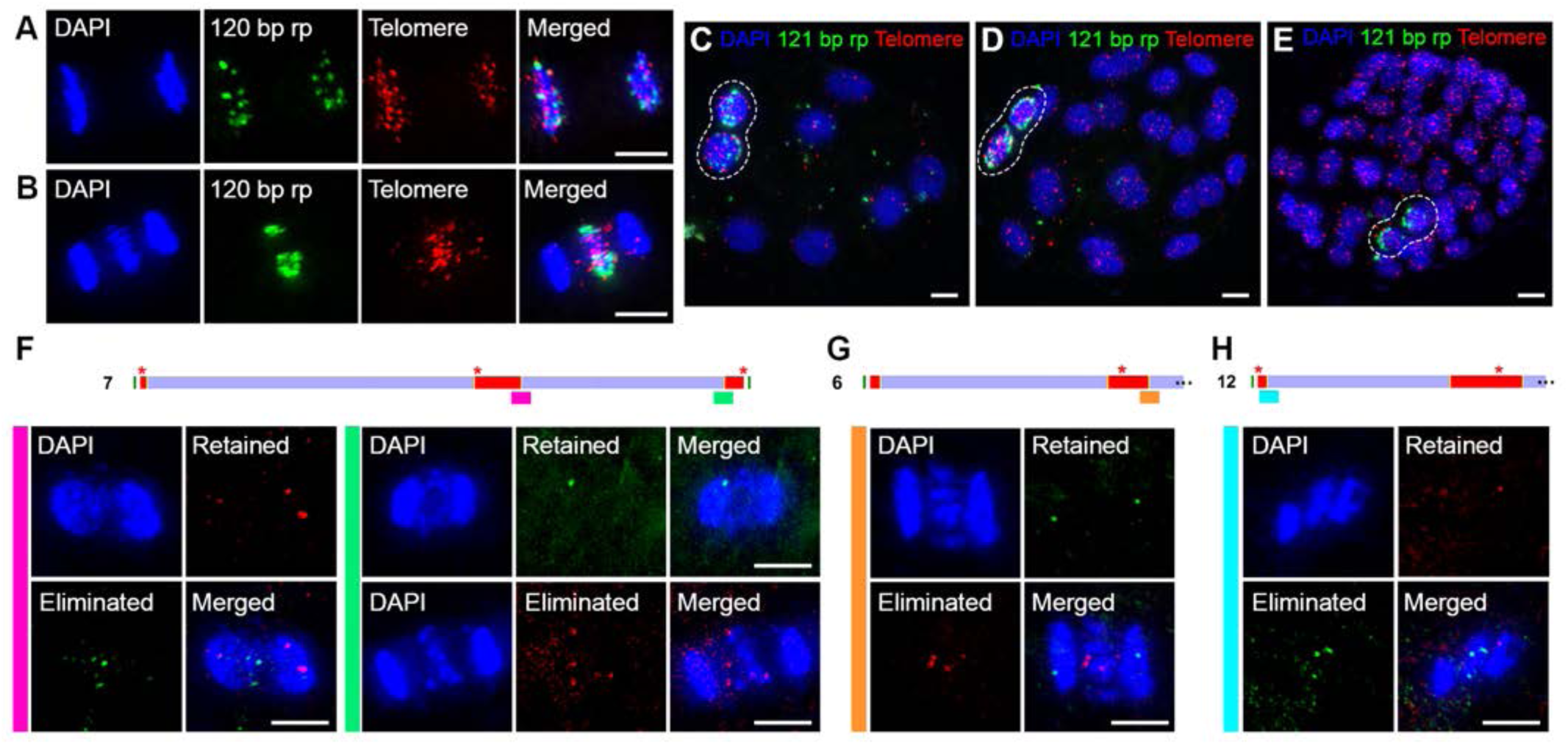
Subtelomeric and telomeric loss in DNA elimination. **(A)** DNA FISH of a non-DNA elimination anaphase in a 3-cell embryo showing that the telomeric and 120 bp tandem repeat DNA are segregated. **(B)** DNA FISH of a DNA elimination anaphase in a 4-cell embryo showing the telomeric and 120 bp repeat DNA remains at the metaphase plate. DNA remaining at the metaphase plate between the segregating DNA is eliminated. **(C)** DNA FISH shows telomeric and 120 bp repeat DNA are not incorporated into daughter nuclei and become cytoplasmic (8-cell embryo). Note the absence of telomeric signal (and 120 bp repeat) in somatic nuclei that have undergone DNA elimination compared to the two germ cells (dotted circle). The new chromosome telomeres following DNA elimination are limited in length. **(D-E)** DNA FISH shows the dynamics of telomere re-extension in somatic nuclei after DNA elimination. **(D)** Post-elimination somatic nuclei in a 24-cell embryo show less telomeric DNA FISH signal with heterogeneous intensities compared to germ cells (dotted circle). **(E)** DNA FISH for telomeres in somatic nuclei of a 64+-cell embryo shows comparable signal intensities to the germ cells (dotted circle) Single channel images for C-E are shown in Supplementary Figure S4. **(F-H)** DNA FISH demonstrates loss of sequences from the ends and internal regions of chromosomes. **(F)** Left panel shows DNA FISH probes corresponding to eliminated (green) and retained regions (red) of an internal chromosome 7 DNA break. Right panel shows DNA FISH probes corresponding to eliminated (red) and retained regions (green) of a subtelomeric chromosome 7 break. **(G)** DNA FISH probes corresponding to eliminated (red) and retained regions (green) of an internal chromosome 6 break. **(H)** DNA FISH probes corresponding to eliminated (green) and retained regions (red) of a subtelomeric chromosome 12 break. Scale bars = 5 µm.

The new genome assembly provided information on the subtelomeric regions of the germline chromosomes that were previously not well-assembled. Comparison of the germline and somatic chromosome termini strikingly demonstrated that CBRs occurred in all subtelomeric regions at both ends of each chromosome (2 x 24 = 48 terminal CBRs in *Ascaris* genome) (Fig. 2 and Table 1). Thus, the data indicate that during DNA elimination all germline chromosome ends undergo subtelomeric breaks, with the loss of distal regions including the telomeres. Our somatic sequencing suggests that the new chromosome ends are healed by *de novo* telomere addition. These data are consistent with the telomere probe DNA FISH data and explain why our molecular analyses did not identify shortened telomeres on the germline chromosomes.

To confirm that the ends of chromosomes were lost during DNA elimination, we carried out DNA FISH using specific subtelomeric probes on each side of the subtelomeric chromosome breaks. As shown in Figure 3F,H, subtelomeric probes corresponding to the eliminated side of a break remain at the metaphase plate during a DNA elimination mitosis and are not undergoing segregation. In contrast, subtelomeric probes corresponding to the retained side of a break are segregated to daughter nuclei. In addition, DNA FISH with probes corresponding to retained and eliminated regions of internal DNA breaks are also consistent with genome data for DNA breaks and sequences lost or retained (Fig. 3F-G). Consistently, in later stage embryos (8 −32+-cells), DNA FISH shows that the eliminated DNA derived from the ends or internal regions of chromosome are absent in somatic nuclei (Supplementary Figure S4). Overall, our DNA FISH and genome comparisons demonstrate that DNA breaks in subtelomeric regions of chromosomes leads to the loss of the all chromosomes ends during *Ascaris* DNA elimination. Thus, *Ascaris* chromosome undergo comprehensive end remodeling during DNA elimination.

### Eliminated DNA

Previous work demonstrated that the eliminated DNA consisted primarily of a tandem 120 bp repeat and unique sequences including 1000 genes [16,22]. The new chromosome assemblies indicate that the eliminated 120 bp tandem repeat constitutes ~ 30 Mb of DNA, 10% of the genome, and 54% of the eliminated DNA. This repetitive sequence is present in 25/48 of the chromosome ends that are eliminated and 9/12 of the internally eliminated regions of chromosome, but absent in all small internal eliminated regions. While the 120 bp tandem repeat is not present in all eliminated regions, genes are present in all regions of the genome eliminated. The size of the eliminated regions containing the 120 bp tandem is a minimum length. The length of the tandem repeat was typically greater than the PacBio reads (on average 900 kb) and thus assembly across the repeat was not possible.

All chromosomes undergo remodeling of the ends of their chromosomes (Fig. 2) with the loss of genes and repetitive sequences from these regions. The range of sequence lost from the ends of chromosomes is from 24 to 829 kb with an average length of 247 kb and a total of 11.5 Mb eliminated. Twelve internal regions of chromosomes are also eliminated (Fig. 2) with a range of sequence lost from 43 kb to 1.6 Mb with an average length of 694 kb and a total of 8.3 Mb eliminated. Thus, while the size of the DNA eliminated from internal regions of chromosomes is on average larger than from the ends of the chromosomes, the total amount of DNA eliminated from the chromosome ends is much greater, particularly when including the 120 bp and subtelomeric repeats lost. Furthermore, while all chromosome ends are remodeled only 11/24 chromosomes (8/19 autosomes and 3/5 sex chromosomes) undergo internal DNA elimination. We estimate the average length of the subtelomeric repeats is ~100 kb and the telomeres are ~15 kb. Overall, the eliminated DNA consists of 30 Mb 120 bp repeat, 11.5 Mb terminal deletion, 8.3 Mb internal deletion, and 5.5 Mb of subtelomeric and telomere repeats. Thus, the total amount of eliminated sequence is 55.3 Mb or 18% of the genome with a germline genome of ~308 Mb.

### Internal CBRs split 11 germline chromosomes into 23 somatic chromosomes

DNA breaks near the ends of all the chromosomes remove the subtelomeric and telomeric sequences, but do not contribute to changes in chromosome number. Twenty-four internal CBRs in 11 chromosomes split the germline chromosomes into 23 smaller somatic chromosomes (Fig. 2 and Table 1) increasing the number of autosomes from 19 in the germline to 27 in somatic cells and the number of sex chromosomes from 5 in the germline to 9 in somatic cells. The largest chromosome (X1) is broken into three somatic chromosomes whereas all other 10 germline chromosomes are split into 2 somatic chromosomes. To determine if there are any features that distinguish the internal vs. terminal CBRs, we carried out DNA motif and other analyses on these two types of CBRs. No distinctive features were identified between these two types of CBRs. In addition, our ATAC-seq data reveals both internal and terminal CBRs becomes equally more chromatin accessible for DNA elimination, with no apparent distinction between these CBRs (see Supplementary Figure S3).

### Histone changes during DNA elimination

During a DNA elimination mitosis, DNA that will be eliminated initially remains at the metaphase plate and is not segregated during anaphase (see Fig. 4A). Following telophase and cytokinesis, this DNA localizes to the cytoplasm where it is eventually degraded (see Fig. 4B). To further examine DNA elimination mitoses and the fate of eliminated DNA, we conducted light and electron microscopy analyses. We first asked whether DNA that was destined for elimination at anaphase of an elimination mitosis exhibited different histone marks compared to DNA that would be retained. Immunohistochemical analysis of several histone marks (H3K4me1/3, H3K9me2/3, H3K36me2/3, H3K16ac, and H3K27me3) during anaphase of a DNA elimination indicated no clear differences between retained vs eliminated DNA (Supplementary Figure S5C). However, H3S10P preferentially marked eliminated DNA (Fig. 4A-B) and H2AK119ub (Supplementary Figure S5B) was reduced in the eliminated DNA during anaphase of a DNA elimination. Histone H3S10P remains on the DNA that is relegated to the cytoplasm. While H3S10P is not present in all DNA fragments in the cytoplasm, this may simply reflect its distribution on chromosomes as H3S10P is not uniformly distributed on chromosomes during prophase of a DNA elimination mitosis (See Supplementary Figure S5A). At telophase of a DNA elimination mitosis, active histone marks (H3K4me3 and H4K16ac, see Fig. 4F-G and Supplementary Figure 5B) are lost on the eliminated DNA. The eliminated DNA in the cytoplasm disappears over time and is gone after several cell divisions. To determine whether autophagy might be involved in the degradation and elimination of the cytoplasmic DNA, we used an antibody against a *C. elegans* marker for autophagosomes LGG-1 [41,42], an ortholog of Atg8/LC3 protein involved in autophagy [43]. LGG-1 IHC staining in *Ascaris* embryos shows distinct LGG-1 puncta, similar to patterns observed in *C. elegans* embryos [41,44,45], and these puncta are coincident with DNA and H3S10P membrane-bound structures (see EM below) suggesting that eliminated DNA degradation likely occurs through autophagy. Overall, our data suggest that the eliminated DNA undergoes dynamic changes over time in both the nucleus and cytoplasm.

**Figure 4.**
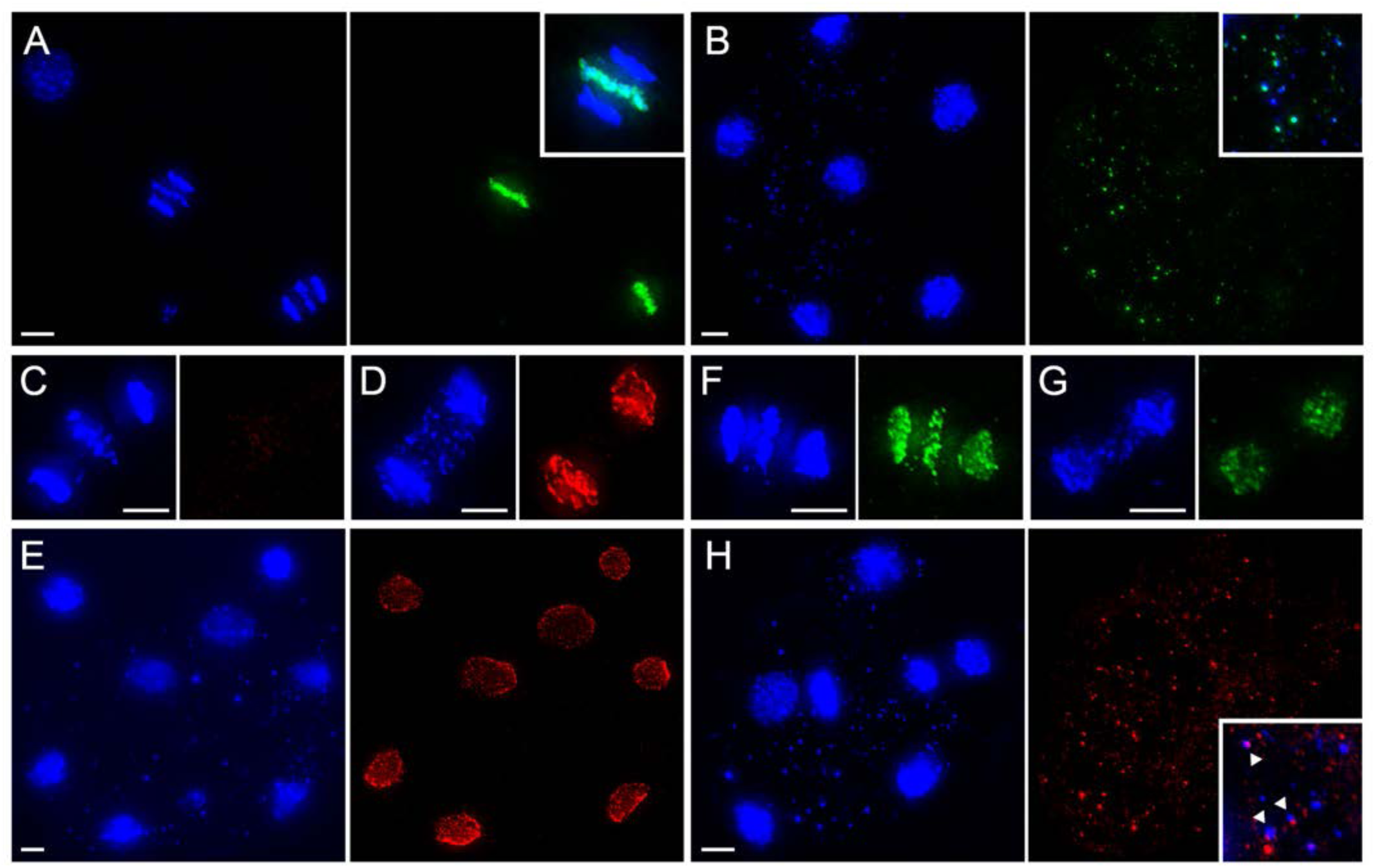
Eliminated DNA is marked with H3S10P, localizes to the cytoplasm, and is associated with autophagosomes. **(A**) Anaphase of 4-cell DNA elimination mitosis. DAPI in blue and H3S10P IHC in green. The fourth cell nucleus is out of the image plane. Inset: Merge DAPI and H3S10P IHC. **(B**) Eliminated DNA fragments in the cytoplasm retain H3S10P. Six cell stage with DAPI DNA fragments in cytoplasm (left panel) and higher magnification of DNA fragments with H3S10P IHC in right panel and in inset. Note that not all DNA fragments in the cytoplasm show H3S10P staining. Chromosomes at prophase prior to a DNA elimination do not exhibit uniform H3S10P staining of chromosomes (see Supplementary Figure S5). **(C-E)** NPC IHC during a DNA elimination telophase. Note that the NPC, absent during anaphase (C), is present in daughter nuclei forming in D and in the nuclei of the seven-cell stage in E, but that eliminated DNA in micronuclei do not exhibit NPCs (see also EM text below). **(F-G)** H3K4me3 IHC shows no differential staining between segregated and eliminated material in DNA elimination anaphase of 4-cell embryo (F) but is depleted from eliminated DNA at telophase (G). **(H)** LGG-1 IHC in a 8-cell embryo suggests DNA containing membrane-bound structures in the cytoplasm form autophagosomes (arrowheads). DAPI staining in left panel, LGG-1 IHC in right panel and inset zoom of merged DAPI and LGG-1 IHC. Scale bars = 5 µm.

**Figure 5.**
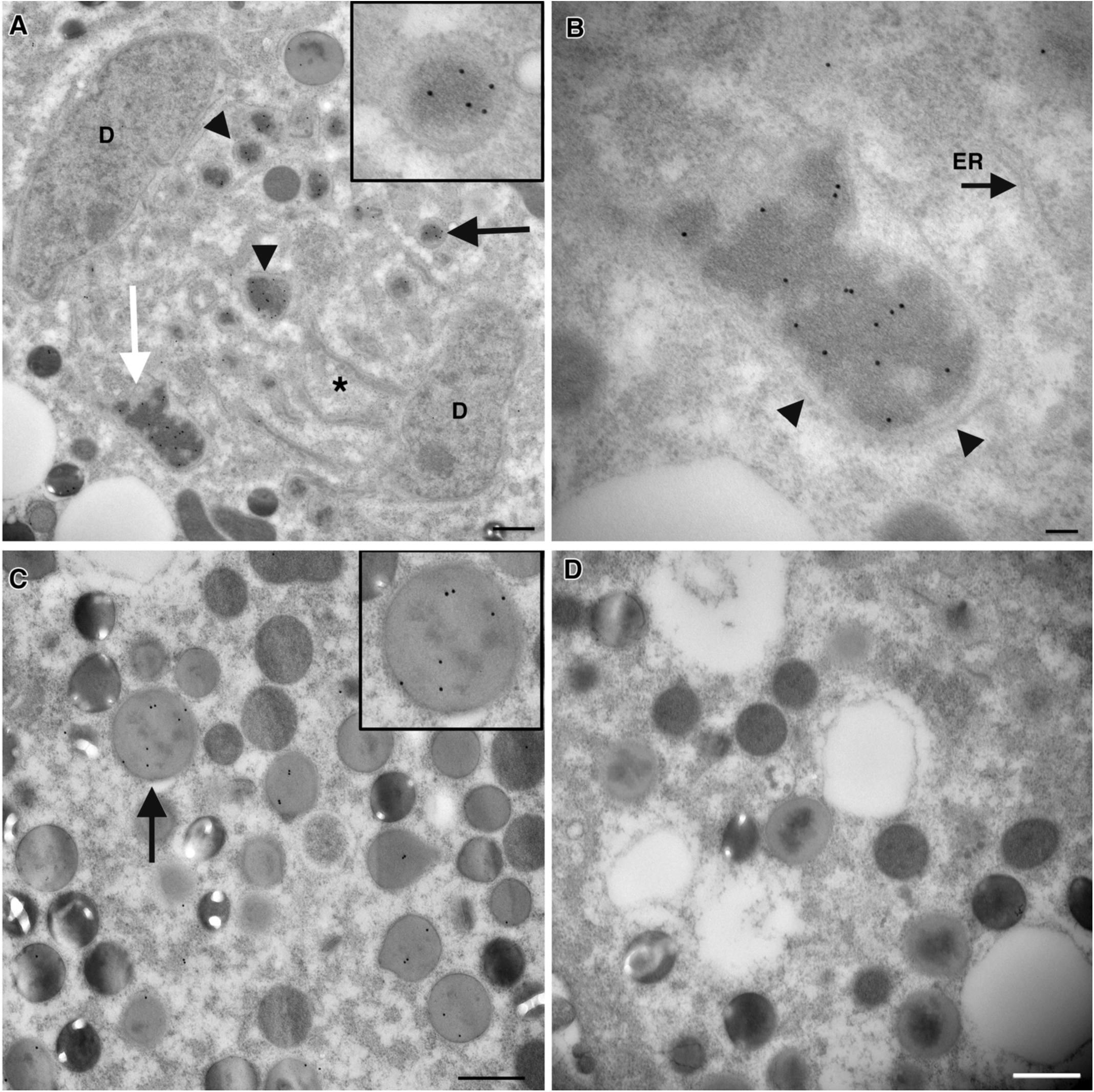
Double membrane-bound structures contain DNA with H3S10P. **(A-D)** Immuno-electron microscopy using an antibody to H3S10P and a secondary antibody coupled to 15nm gold. **(A)** Telophase cell in a DNA elimination mitosis. Numerous membrane-bound organelles containing electron dense material label with this antibody (arrowheads; black arrow points to granule surrounded by double membrane in inset). Bar= 500nm. Membrane compartments containing dense material (above and below the star) were also observed spanning the region of DNA elimination. (B) H3S10P antibody preferentially labels dense material within double membrane-bound organelles (black arrowheads). ER, endoplasmic reticulum. Bar = 100nm. (**C)** H3S10P antibody labels numerous structures in the cytoplasm of post-diminution cells. Bar = 500nm. **(D)** Control samples with the secondary antibody-gold show no labeling. Bars = 500 nm.

### Eliminated DNA is packaged into double membrane-bound structures (micronuclei)

Electron microscopy of DNA elimination mitoses identified numerous, double-membrane bound structures of heterogeneous size and shape in telophase cells (Fig. 5A-B). These structures appear when the nuclear membrane reforms around the retained daughter nuclear DNA and are present in the cytoplasm of cells that have undergone DNA elimination. Immuno-EM (using an antibody to H3S10P) demonstrated that the telophase structures and those present in the cytoplasm following DNA elimination were positive for H3S10P, indicating the presence of chromatin. These results indicate that the eliminated DNA is compartmentalized into double-membrane bound structures (Fig. 5A-B; insets). Overall, the structures have characteristics of micronuclei and will hereafter be called micronuclei [46]. The H3S10P antibody preferentially labeled electron-dense material within the core of the structures, indicating the material is hyper-condensed DNA (Fig. 5B). In addition, the cytoplasm of cells after DNA elimination contained numerous structures that labeled with the H3S10 antibody (Fig. 5C). Serial section immuno-EM confirmed labeling on the same granules appearing in adjacent sections (data not shown). No labeling was detected on control sections (Fig. 5D). Consistent with what was observed by IHC, the H3S10P antibody did not label DNA that had segregated to daughter nuclei (see Fig. 4, Fig. 5A). Notably, based on IHC and EM, the micronuclei have limited or no nuclear pores (see Fig. 4C-E; Fig. 5).

### Ultrastructural timeline of DNA elimination

We next used electron microscopy to study the timeline of DNA elimination. Serial semithick (200nm) sections were imaged to identify cells at various mitotic elimination stages. Late prometaphase spindles contain highly condensed chromosomes near the metaphase plate (Fig. 6A). Surprisingly, lighter staining, more diffuse chromatin was observed near the metaphase plate and in the body of each half-spindle (Fig. 6A, *’s). Electron tomography of adjacent sections was performed to characterize the 3D organization of spindle MTs, both at the condensed chromosomes and the lighter stained, more diffuse chromatin (Fig. 6A, inset model). Shown in this model and in Movie S1, a total of 460 MTs were tracked and 115 MTs were identified as kinetochore microtubules KMTs that made end-on attachments to kinetochores on condensed chromosomes (Fig. 6A; purple; Movie S1). Such structures and connections were only found on condensed chromosomes. The other 345 mts tracked (Fig. 6A, green; Movie 1) were identified as non-KMTs; these either did not end on chromosomes or went out of the volume of the reconstruction. These non-KMTs surround the regions containing diffuse chromatin but do not make end-on attachments (Movie S1). The diffuse chromatin in the half-spindles was structurally different from the highly condensed chromosomes that lay predominantly at the metaphase plate (compare Fig. 6B and 6C) and appeared more lightly stained and less compact. Tomographic reconstructions revealed that this material does not assemble kinetochores and does not contain MT end-on attachments. Instead, MTs pass this material making close, lateral contacts (Fig. 6B, arrowheads; Movie S2). In contrast, the highly condensed chromosomes at the metaphase plate appear more compact and assemble clear kinetochores to which KMTs attach (Fig. 6C). Supplementary Figure S6 and Movie S3 show that even when diffuse chromatin was situated near condensed chromosomes, only the condensed chromosomes assemble kinetochores and have KMTs. A larger tomographic volume was computed from a different region of the prometaphase spindle. Of the 663 MTs tracked, 187 were identified as KMTs (Fig. 6D, Supplementary Figure S6C-D; purple; Movie S4) that were only at the condensed chromosomes. Again, there were no recognizable kinetochores or end-on attachments of MTs (green; n=476) to the diffuse, less condensed chromatin present in the volume (Figure 5G, green; Movie 4). These were found only at the condensed chromosomes. These data demonstrate that there are two distinguishable forms of chromatin condensation, one of which leads to normal chromosomes whereas the other leads to chromatin masses that do not assemble kinetochores or bind to kMTs. This diffuse, less condensed chromatin that will be eliminated can be distinguished ultrastructurally from the condensed chromosomes and can be identified in the spindle prior to metaphase.

**Figure 6.**
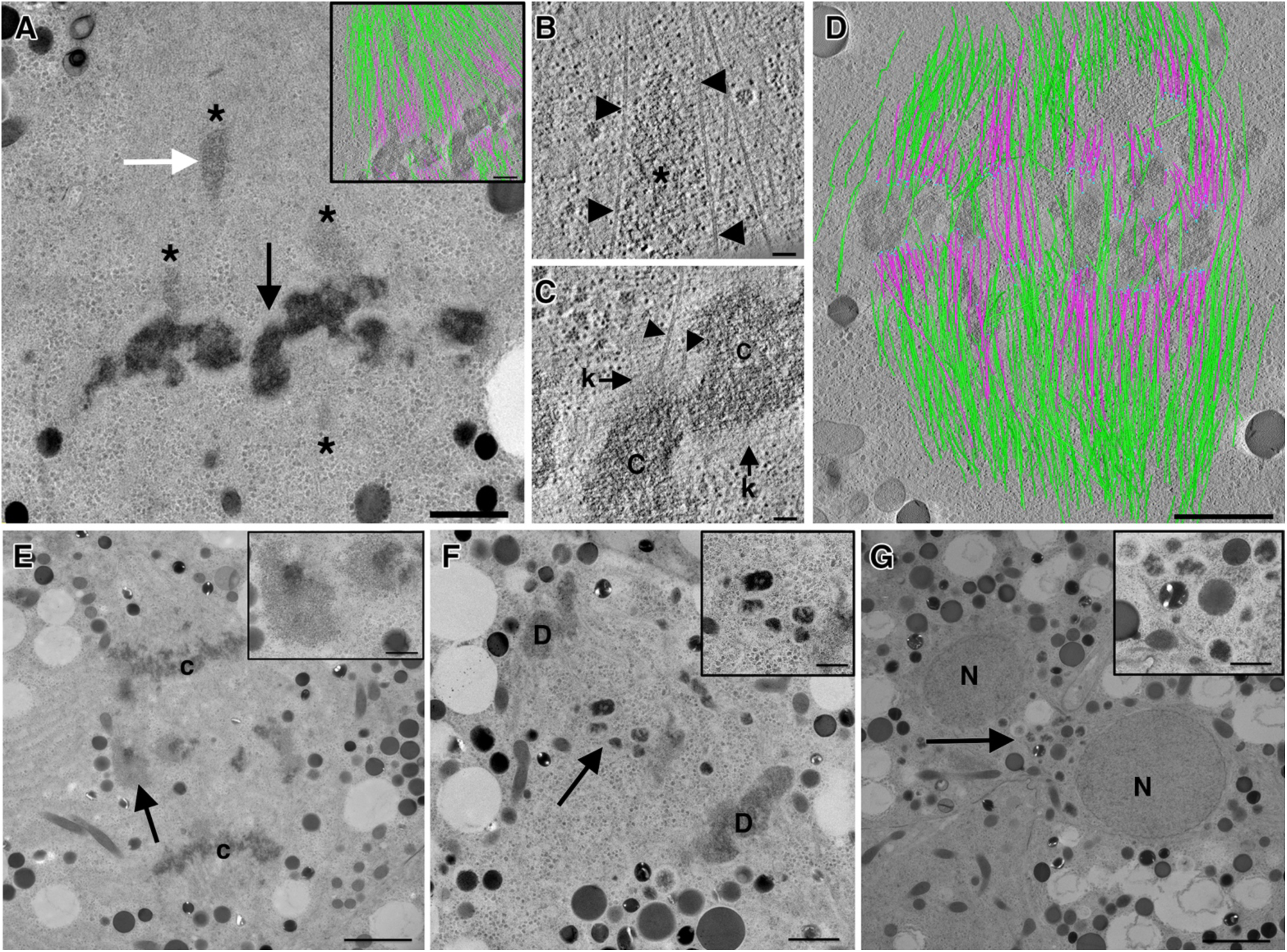
Electron microscopy provides an ultrastructural timeline of *Ascaris* DNA elimination. **(A-D)** Late prometaphase of a DNA elimination mitosis at the 4-cell stage. **(A**) Overview shows darkly stained, condensed chromosomes at the metaphase plate. Lighter staining chromatin (*’s) was observed in the spindle proper and close to the metaphase plate. Bar = 1 µm. Inset shows a model from a tomographic volume computed from two serial sections (green, MTs that do not end on chromatin or go out of the reconstructed volume; purple, KMTs that end on the chromosomes). Serial, tomographic slices and projected 3D model from this region are shown in Movie S1. White and black arrows indicate regions from selected tomographic slices shown in **B** and **C**, respectively. **(B)** Tomographic slice showing fine structure of the lighter staining, diffuse chromatin in the spindle (white arrow in A). Kinetochores were not observed on this diffuse chromatin but it was always surrounded by numerous MTs in close, lateral proximity (arrowheads; bar = 100 nm). Serial, tomographic slices showing modeled MTs in this region are shown in Movie S2. **C)** In contrast, the highly condensed chromosomes (**C)** at the metaphase plate assemble kinetochores (k) which are associated with KMTs (arrowheads) Bar = 100nm. Movie S3 shows serial, tomographic slices from another area showing KMTs at chromosomes but not at the diffuse chromatin. **(D)** 3D model of the MTs in 3 serial, 200nm thick sections from the prometaphase spindle. KMTs (purple) end on condensed chromosomes only. Bar = 1 µm. Serial, tomographic slices and projected 3D model from this region are shown in Movie 4. **(E-G)** Double membranes begin to surround eliminated DNA at telophase. (**E)** Overview of a DNA elimination mitosis in anaphase shows segregated chromosomes (C) at each pole with eliminated DNA at the spindle equator. Arrow points to mass of lightly condensed material. Higher magnification (inset) shows that this material is not surrounded by membrane. Bar = 2 µm; inset bar= 500nm. **(F)** Membranes begin to surround eliminated DNA at telophase. Nuclear envelope begins to form around segregated DNA at the spindle poles (D’s). Numerous membrane compartments are observed throughout region of the spindle midplane (*). Granules containing eliminated DNA appear more darkly stained and are surrounded by double membranes (arrow, inset) Bar= 500nm; inset bar= 500nm. **(G)** Cytokinesis following a DNA elimination mitosis shows two daughter nuclei (N) and numerous electron-dense, membrane-bound granules near the spindle midbody (arrow, inset). Bar=2 µm; inset bar= 500nm.

EM of DNA elimination anaphase spindles show segregated chromosomes near the spindle poles and eliminated DNA remaining at the spindle midplane (Fig. 6E; c’s and arrow, respectively). MTs were observed connecting the segregated chromosomes to the centrosomes of each pole (data not shown). The eliminated DNA at the spindle midplane was not attached to MTs and appeared as masses of diffuse chromatin similar in structure to that observed in prometaphase (Fig. 6E; inset). This material was not surrounded by membrane at this stage. The nuclear envelope begins to reform around segregated DNA at late anaphase/telophase (Figs. 5A, 6F). The telophase spindle contained numerous membrane-bound compartments in the region spanning the spindle midplane (Figs. 5A and 6F; *), some of which contain electron dense material (Fig. 5A). Double membranes surrounding the eliminated DNA, micronuclei, are first observed at this stage (Fig. 6F, inset; 5A). The DNA in these structures becomes more condensed compared to the diffuse chromatin observed in prometaphase and anaphase. At cytokinesis, the micronuclei are observed near the cleavage furrow (Fig. 6G, arrow). Overall, the EM data suggest that retained and eliminated DNA can be identified and differentiated with the diffuse, less condensed chromatin, DNA for elimination, adjacent to the metaphase plate and without kinetochores and kMTs. DNA FISH for the eliminated 120 bp repeat (see Supplementary Figure S7) also shows that this eliminated DNA is adjacent to the metaphase plate.

### Chromosome fusion in *Ascaris* sex chromosomes

*Ascaris* has a large number of germline chromosomes compared to most other known nematodes (most have 4-12 or less) [47]. To further examine *Ascaris* chromosomes, particular the unusual multiple sex chromosomes, we compared the 24 *Ascaris* germline chromosomes to the 6 chromosomes in the free-living nematode *C. elegans* which likely represents an ancestral chromosome state (Fig. 7). All 19 *Ascaris* autosomes can be traced to an individual *C. elegans* chromosome; thus, the expansion of *Ascaris* autosomes is likely due to the subdivision of individual ancestral chromosomes (Fig. 7). While in general gene linkage for these 19 autosomes is maintained between the two species, there is little overall chromosomal synteny. This suggests that there has been extensive genome rearrangement within chromosomes after the separation of the species, as seen in other nematodes [48–50]. In contrast, the majority of *Ascaris* sex chromosomes (X1, X2, X4 and X5) were traced to multiple *C. elegans* chromosomes (Fig. 7A-B). While some rearrangements are present in chromosome X5, the three other *Ascaris* sex chromosomes (X1, X2 and X4) are devoid of rearrangement between the blocks of chromosomes, particular for the blocks matching *C. elegans* chromosome I. This suggests that these *Ascaris* sex chromosomes are likely derived from recent chromosome fusion events. Interestingly, the boundaries for these fused chromosome blocks coincide with the *Ascaris* CBRs (Fig. 7B). With the DNA breaks at the CBRs during DNA elimination, the *Ascaris* sex chromosomes become orthologous to single *C. elegans* chromosomes following DNA elimination. Finally, the *Ascaris* eliminated DNA regions have very few unique orthologous genes to *C. elegans* (Fig. 7A-B), suggesting that these eliminated regions have recently evolved or are evolutionarily more flexible than the rest of the genome. These regions may more readily enable the birth of new genes, consistent with the observation that novel genes are often enriched in the testis and consistent with their propensity for elimination in *Ascaris* [51–53]. Overall, our comparative analysis of the *C. elegans* and *Ascaris* chromosomes reveals that many *Ascaris* sex chromosomes are the result of recent fusion events and that DNA elimination breaks re-subdivide the chromosomes into single chromosomes that are orthologous to individual *C. elegans* chromosomes.

**Figure 7.**
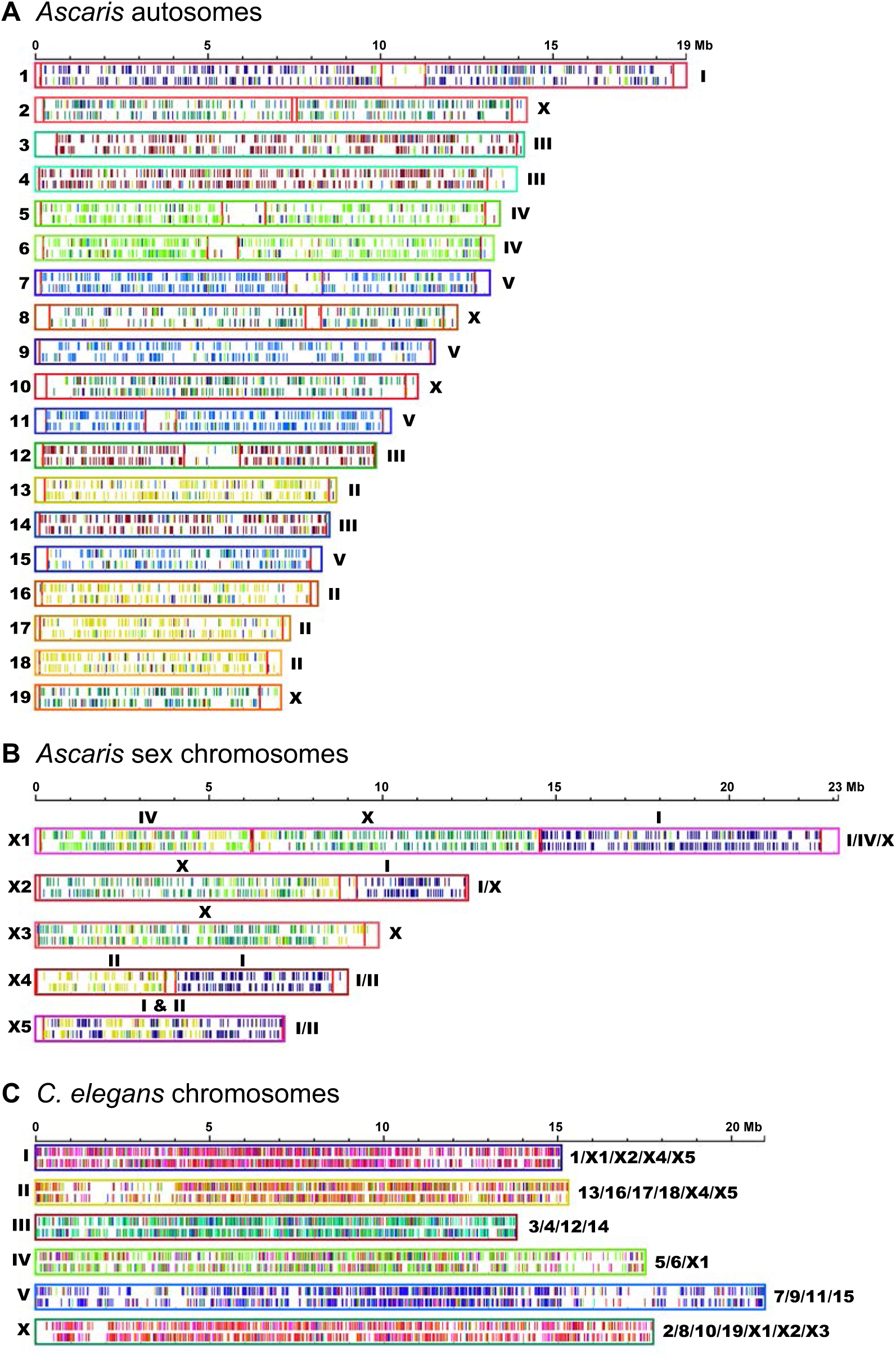
Comparison of *Ascaris* and *C. elegans* chromosomes. The reciprocal best blast hit for *Ascaris* and *C. elegans* orthologous proteins (6,402) were identified, the genes encoding these orthologous proteins shown as vertical half lines with the color illustrating the origin in the orthologous nematode chromosomes (top half indicates forward strand). The chromosome outer box color corresponds to the color of the vertical lines for the orthologous genes in the other nematode. The chromosomes are labeled on the left and their corresponding major chromosomes in either *C. elegans* or *Ascaris* labeled on the right (and also on the top for *Ascaris* sex chromosomes). Vertical red lines in the Ascaris chromosomes indicate the chromosomal break regions where DNA breaks occur during DNA elimination.

## Discussion

We describe significantly improved germline and somatic genome assemblies for the parasitic nematode *Ascaris* that provides novel insights into programmed DNA elimination. The full-chromosome assemblies provide the first and most comprehensive chromosomal view of DNA elimination in a metazoan. Comparison of these assemblies demonstrated that all germline chromosomes undergo breaks in subtelomeric regions with loss of distal sequences and healing of chromosome ends by *de novo* telomere addition. Overall, we have identified 72 chromosome breaks (32 new CBRs in this study). Terminal chromosome breaks remodel all 48 chromosome ends while 24 internal DNA breaks (11/24 chromosomes) increase the somatic chromosome number from 24 to 36. EM and IHC analyses indicate that eliminated DNA is packaged into double membrane-bound structures, which we call micronuclei, at telophase. These micronuclei undergo chromatin changes over time including the loss of active histone marks. At the end of a DNA elimination mitosis, the micronuclei localize to the cytoplasm, form autophagosomes, and the DNA is eventually lost. Comparative analysis of the *Ascaris* chromosomes suggests that *Ascaris* sex chromosomes were formed by chromosome fusions that are resolved into independent chromosomes through the DNA breaks at CBRs during DNA elimination. Our comprehensive genomic and cytological studies on DNA elimination provide key new insights on metazoan DNA elimination.

### Remodeling of chromosome ends

Using DNA FISH for telomeres, a previous [30] and our study observed that telomeres appeared to be removed from all *Ascaris* chromosomes during a DNA elimination and subsequent telomere length extension was asynchronous requiring several cell cycles (see Fig. 3 and Supplementary Figure S4). Our genome analyses indicate that during DNA elimination subtelomeric chromosome breaks occur releasing the distal portions of chromosome with their intact telomeres. DNA FISH experiments support that chromosomes undergo subtelomeric breaks and confirm the loss of distal chromosome regions and their telomeres during DNA elimination. Thus, all *Ascaris* germline chromosomes undergo chromosome end remodeling during DNA elimination (see Fig. 2, 3, and Supplementary Figure S4).

The new finding that all germline chromosome ends undergo remodeling through subtelomeric breaks and addition of *de novo* telomeres raises a variety of questions. For example, what impact does the remodeling have on chromosome architecture. Subdivision of single chromosomes into 2 or more chromosomes and the remodeling of the chromosome ends is likely to alter the overall architecture of each chromosome and inter-chromosome interactions. It remains to be determined what overall impact DNA elimination has on *Ascaris* chromosome architecture and whether chromosome architecture prior and during DNA elimination might contribute to DNA elimination as recently observed in the ciliate *Tetrahymena* [54]. An open question is whether the breaks in subtelomeric regions (and internal regions) are facilitated by chromosome looping mechanisms that contribute to the DNA breaks. Further high-resolution and stage-specific Hi-C and FISH work is needed to address these questions.

The remodeling of chromosome ends may also impact the overall gene expression of the chromosomes. Muller [55] first suggested that the chromosome breaks and addition of new telomeres may lead to a “telomere position effect” on gene expression following DNA elimination [56]. While previous studies on a single CBR [55] and within 50 kb of a number of CBRs [16] have been carried out using steady-state RNA levels analyses, no pattern of gene silencing or activation were identified. The current study identifies significant additional remodeling of chromosome ends. Future comprehensive nascent RNA sequencing before, during, and after DNA elimination is required to determine if chromosome end-remodeling impacts gene expression near the ends of the newly remodeled chromosomes.

### DNA Elimination of the *Ascaris* 120 bp repeat

The majority of the eliminated DNA (54%) consists of a 120 bp tandem repeat, with over 99% of this repeat lost. The large amount of repeat eliminated has historically led to the speculation that the repeat may be mechanistically involved in the elimination process. Our new chromosome assemblies enabled us to identify the location of these repeats on the chromosomes and their proximity to the DNA breaks. The location of the 120 bp tandem repeats within the eliminated regions is variable, but typically not near where the chromosomal DNA breaks (CBRs) occur; the majority of the CBRs (50/72, 70%) are not within 200 kb of the repeats. This suggests it is unlikely the 120 bp repeats play a direct role in the DNA breaks and DNA elimination. However, we cannot rule out that the repeats may contribute to some higher order 3D chromosome organization that contributes to DNA elimination. Notably, not all eliminated genome regions contain the 120 bp tandem repeat. However, all eliminated genome regions contain genes. This indicates that the loss of genes is not simply a consequence of being linked to regions of the genome where the 120 bp tandem repeat is located. Thus, *Ascaris* DNA elimination serves not to just simply remove the 120 bp repeats. The findings support our model that the elimination of genes is a key component of DNA elimination [16,22].

### DNA elimination in other organisms

In ciliates, formation of macronuclear chromosomes (somatic genome) results from many chromosome breaks in the 5 micronuclear chromosomes (germline) and the healing of the broken ends by telomere addition. In *Tetrahymena thermophila*, for example, 225 chromosome break sites leads to 181 small amplified (~45 copies on average 900 kb chromosomes) macronuclear chromosomes (somatic) [57]. The chromosome end remodeling observed in *Ascaris* differs from that described in ciliates in that the breaks do not generate a large number of relatively small chromosomes and no amplification occurs.

In copepods, large amounts of DNA are eliminated while chromosome numbers, in general, do not change [8,58]. One model for DNA elimination in copepods is that internal elimination within chromosomes involves looping out of DNA, excision of the DNA, and ligation of the ends. However, the contribution of internal vs terminal DNA elimination remains unknown. Programmed DNA elimination in lampreys is associated with decreased chromosome numbers (chromosome loss and/or fusion) and loss of ~20% the germline DNA (500 Mb of DNA) in somatic cells, loss of both repetitive sequences and some germline expressed genes [11,12,14,15]. In hagfish, loss of only repetitive sequences has been identified in DNA elimination [9,10]. DNA FISH and cytogenetic analysis indicate the loss of whole hagfish chromosomes and chromosome termini encompassing 21-75% of the germline DNA in different species [9,10]. It is not known whether the loss at hagfish chromosome termini is similar to that observed here in *Ascaris* or if an excision event (internal deletion and fusion) occurs removing subtelomeric regions without loss of the telomere from the chromosome. Overall, the current studies are the most comprehensive genome analysis of DNA elimination.

### DNA breaks

Analysis of all the sites for DNA breaks (including 32 new sites in the current study), further supports that there are no sequence, structural motifs, histone marks, small RNAs, or other features that define or target the 3-6 kb CBRs as sites for the DNA breaks in chromosomes [16,22–25]. Overall, these characteristics suggest it is unlikely a sequence specific endonuclease is involved in generating the DNA breaks. However, we cannot rule out that an enzyme similar to a Type I restriction enzyme [59] that recognizes unique sequences at a distance from the DNA breaks might be involved in generating the breaks randomly within the CBRs.

Domesticated transposases, such as RAG1-RAG2 for V(D)J recombination [60–62] and PiggyBac in ciliate DNA elimination [63–66], recognize specific sequences in the genome during development to generate DSBs. However, we have not identified any transposases or their signatures in *Ascaris* that might generate the DNA breaks. In ciliates, programmed DNA rearrangements involves the elimination of short internal sequences followed by re-ligation of the boundaries (IES) and many chromosomal break sites (CBS) that lead to new chromosomes and loss of sequences [17–21]. While there is some understanding of the breaks at IESs, the mechanism for the CBS is unknown. Thus, the mechanism(s) for key DNA breaks in DNA elimination remain unknown. Open chromatin previously identified likely contributes to identification of regions for the *Ascaris* breaks, facilitates the generation of the breaks, and/or is a consequence of molecular processes that contribute to the generation of the breaks [16].

### Compartmentalization of eliminated DNA into micronuclei during telophase

Our EM studies using high pressure freezing and freeze substitution methods [67–69], leading to optimum embryo preservation, provide an ultrastructural timeline of *Ascaris* DNA elimination. We observed differential chromatin condensation in late prometaphase spindles, as previously observed using light microscopy [30]. Our EM images showed that there are two distinguishable forms of chromatin condensation, one leads to normal chromosomes whereas the other leads to chromatin masses that are less diffuse and compact. The diffuse, less condensed chromatin destined for elimination can be distinguished ultrastructurally from the condensed chromosomes and can be identified near the metaphase plate and in the spindle proper in late prometaphase. In addition, electron tomography was used to study the 3D arrangement of spindle MTs, with particular focus on the arrangement of individual MTs around the diffuse chromatin in the prometaphase spindle. Electron tomography confirmed that this material does not assemble a kinetochore or make end-on connections with KMTs; it is instead surrounded by lateral MTs. This is consistent with the fact that CENPA is significantly reduced in chromosome regions that will be lost during *Ascaris* DNA elimination [28] (this study) and EM studies in *Parascaris* that showed chromatin that will be eliminated does not assemble kinetochores or bind KMTs [27]. Notably, DNA FISH (see Supplementary Figure S7) indicates that the eliminated 120 bp repeat is present at the boundary of the metaphase plate and appears to be similar to the diffuse, less condensed observed in EM. Furthermore, previous studies suggested that chromosome DNA destined for elimination in both *Ascaris* and *Parascaris* was less condensed [30]. Overall, these data show that DNA for elimination undergoes differential condensation and is often situated at the boundary of the metaphase plate. The diffuse chromatin is not segregated in anaphase, remaining in the spindle midplane as retained chromosomes move to the poles. Concurrent with nuclear envelope reformation at telophase, the eliminated DNA is surrounded by a double membrane at telophase forming structures similar to micronuclei. During telophase and cytokinesis, the DNA within the micronuclei becomes more condensed which may be related to the loss of active histone marks. These micronuclei persist in the cytoplasm following the end of a DNA elimination mitosis for some time and eventually disappear. The EM studies presented thus provide new details of mitotic spindle organization and further our understanding of the ultrastructural timeline of DNA elimination in Ascaris.

Immunohistochemistry studies demonstrated that DNA which remained at the metaphase plate and is eliminated contained high levels of histone H3S10P whereas segregated and retained DNA did not (see Figs. 4-5). We interpret the retention of H3S10P on DNA that will be eliminated to indicate that this DNA does not undergo the metaphase to anaphase transition. Notably, histone H3S10P remains on eliminated DNA in the cytoplasm until the DNA is eventually degraded. Immuno-EM for H3S10P demonstrates that material in the double membrane-bound vesicles during telophase and in the cytoplasm contains DNA (see Fig. 5). DNA FISH on the cytoplasmic material also demonstrates that the micronuclei contain eliminated DNA (see Fig. 3). Thus, eliminated DNA is sequestered into micronuclei at telophase during a DNA elimination and chromatin undergoes changes in histone modifications. The micronuclei have significantly reduced nuclear pore complexes (Fig. 4-5).

Micronuclei have been described associated with DNA elimination in lampreys, the cartilaginous rat fish, and water frogs [6,13,14]. EM studies in water frogs and the rat fish provide structural evidence for micronuclei demonstrating double membrane bound vesicles. Micronuclei in water frogs exhibited less nuclear pore complexes and greater heterochromatization than the main nuclei. In lampreys, micronuclei did not possess lamin B1 nor nuclear pore O-linked glycoproteins. Thus, eliminated DNA appears to be incorporated into micronuclei-like vesicles in all these organisms. The micronuclei-like vesicles in the cytoplasm of *Ascaris* embryos show association with LGG-1 puncta suggestive of different stages of autophagosome formation (see Fig. 4H). Similar associations have been observed in DNA elimination events in water frogs [6] and the ciliate *Tetrahymena [70], a*nd autophagy has been shown to play a role following meiosis in the programmed nuclear death of the old macronucleus in *Tetrahymena* In the copepod, *Cyclops kolensis*, eliminated DNA is present in double membrane-bound structures in late interphase and early prophase of an elimination division [54]. We recently observed similar double membrane-bound structures in another copepod, *Mesocyclops edax* (Zagoskin, M, O’Toole, E., and R.E. Davis, unpublished data). In *C. kolensis*, the double membrane-bound structures increase in size, speculated to result from fusion of the structures, and the double membrane is reduced to a poreless single membrane by late telophase. These structures with the DNA become cytoplasmic and are rapidly degraded.

All DNA fragments that result from DNA breaks are healed with telomeres [71]. However, when they are healed remains unknown. Furthermore, free DNA ends are problematic in cells leading to possible fusions and recombination. The sequestration of eliminated DNA into vesicles may be involved in both its compartmentalization to avoid these consequences as well as a mechanism to target the DNA for degradation.

### Timing of DNA breaks

Previous models for Ascaris DNA elimination postulated that the DNA breaks likely occurred at the metaphase plate during an elimination mitosis [30]. Our previous data indicate that chromosome regions that will be lost have reduced CENP-A and kinetochores prior to a DNA elimination mitosis [28]. If the DNA breaks occurred prior to mitosis, some alternative mechanism would be required to congress eliminated chromosome fragments to the metaphase plate. The EM structural data described here suggest there are chromosome fragments at the boundary of the metaphase plate during a DNA elimination mitosis that do not have kinetochores or kMTs attached. This suggests that DNA breaks in the chromosomes may have occurred prior to metaphase. In addition, DNA FISH for the eliminated 120 bp repeat indicates this DNA localizes adjacent to the metaphase plate during a DNA elimination. Thus, these data suggest that the DNA breaks in DNA elimination likely occur prior to mitosis. EM observations clearly indicate that the eliminated DNA must reach the metaphase plate through mechanisms that are independent of kinetochores and kMTs. Thus, alternative mechanisms for congression of the eliminated DNA to the metaphase plate must be involved since the DNA lacks CENP-A and kinetochores [28]. These mechanisms could include the contribution of polar ejection forces involving dynamic instability of pole-initiated MTs, plus-end directed chromokinesins, the contribution of interpolar microtubules, or scaffolding or tethering proteins within the spindle [72–77].

### Ascaris sex chromosome fusions and their separation in somatic cells by DNA elimination

Nematodes typically have a small number of chromosomes (4-12)[47]. Assuming that the free-living *C. elegans* is an ancestral chromosome state with six chromosomes, our chromosome comparisons indicated that all of the *Ascaris* autosomes likely represent independent regions of these ancestral chromosomes without fusions (see Fig. 7). In contrast, several *Ascaris* germline sex chromosomes appear to be fusions of independent regions of *C. elegans* chromosomes (see Fig. 7B; for example, *Ascaris* X1 = fusion of regions of *C. elegans* I, IV, and X; *Ascaris* X2 = fusion of regions of *C. elegans* I and X; and *Ascaris* X4 = fusion of regions of *C. elegans* I and II). These fused regions become independent chromosomes in somatic cells after DNA elimination as the regions are separated by CBRs that break the chromosomes. Extreme fusion of chromosomes appears to have occurred in some ascarids where the complete germline genome becomes a single chromosome as in *Parascaris univalens*. Notably, following DNA elimination, this single *P. univalens* germline chromosomes is subdivided into 35 somatic chromosomes [30]. Thus, while the fusion of chromosomes may be accommodated in the germline genome, their separation by DNA elimination chromosome breaks may be necessary in somatic cells. Overall, we suggest that *Ascaris* ancestral chromosomes may have undergone multiple breaks to form many smaller chromosomes. These chromosomes fused to form multiple germline sex chromosomes, only to be separated in somatic cells during DNA elimination. An open question is why these sex chromosomes must be independent chromosomes in somatic cells (1 germline to 6 somatic in *Parascaris* and 5 germline to 9 somatic in *Ascaris*).

## Conclusions

Programmed DNA elimination occurs in diverse organisms from ciliates to vertebrates. Here, we have generated chromosome assemblies of all *Ascaris* germline and somatic chromosomes. Comparison of these assemblies provides the first chromosome view of metazoan DNA elimination including the location of all DNA breaks and the locations of eliminated sequences including repeats and genes. The predominant DNA elimination events in *Ascaris* are comprehensive chromosome end remodeling through breaks in the subtelomeres of all 24 germline chromosomes, loss of distal chromosome sequences, and healing of the breaks by de novo telomere addition. The major eliminated sequence, a 120 bp satellite repeat (30 Mb), is not present in all eliminated regions of the genome. All eliminated DNA contains genes (1000 genes in total) that are predominantly expressed in the germline. We provide an ultrastructural timeline for a DNA elimination mitosis and describe the incorporation of eliminated DNA into double membrane-bound structures, micronuclei, at telophase. The micronuclei exhibit loss of active histone marks and appear to be degraded by autophagy. Our data also suggest recent chromosome fusions in the germline that are broken into multiple sex chromosomes during DNA elimination. These studies provide the most comprehensive genome analysis of DNA elimination in a metazoan, an ultrastructural analysis of DNA elimination, and identify additional novel and key aspects of metazoan DNA elimination.

## Materials and Methods

### *Ascaris* and megabase-size DNA isolation

Collection of *Ascaris* tissues, zygotes, and embryonation were as previously described [36,78]. DNA isolation for PacBio sequencing *(and Southern Blots)* was prepared from embryos embedded in agarose. The chitinous shell was removed by treatment for 90 min in 0.5 N NaOH and sodium hypochlorite at 37°C followed by extensive washing in PBS, pH 7.4. The embryos were treated for 1 min in 90% isopropanol (this removes the ascaroside-based impermeable layer), washed with PBS, pH 7.4, suspended in 10 mM Tris, pH 7.2, 20 mM NaCl, 50 mM EDTA and embedded in 2% CleanCut ™agarose (BioRad) in BioRad plug molds for a final agarose concentration of 0.75%. Agarose plugs were incubated in 2.5 ml of lysis buffer (10 mM Tris pH 8, 25 mM EDTA, 100 mM NaCl, 0.5% SDS) with 1.4 mg/ml Proteinase K (Ambion, AM2548) in a 50°C water bath for 2 hr with intermittent mixing and then overnight with fresh Proteinase K solution. The plugs were extensively rinsed and washed (3X rinse and 5X wash) in 10 mM Tris pH 8.0, 50 mM EDTA and then rinsed and washed twice in 10 mM Tris pH 8.0, 1mM EDTA). To extract genomic DNA, the plugs were digested with beta-agarase (NEB M0392) as described by the manufacturer to liberate DNA, followed by drop-dialysis (VCWP04700 Millipore) against 15 ml of TE buffer. The DNA was further treated with 0.1% SDS and 80 μg of proteinase K *(need concentration here)* before precipitation with 15 µg of glycoblue carrier (Ambion, AM9515), 0.3 M NaOAc pH 5.2, and 2.5 volumes of 100% ethanol. The precipitated DNA was dissolved in low TE buffer (10 mM Tris-HCl pH 8.0, 0.1 mM EDTA) or water.

### PacBio sequencing and initial assembly

Two PacBio libraries derived from 2-cell (48hr) and 4-cell (65hr) embryo DNA were prepared and sequenced by the University of Washington PacBio Sequencing Services (https://pacbio.gs.washington.edu/) using the Sequel II system. From two SMRT Cell runs, we obtained ~360X raw reads with average read length of 15 kb. PacBio reads over 40 kb (average 52 kb, total ~100X read coverage) were used to assemble the initial germline genome assembly using Canu [29] with parameters “corOutCoverage=1000 correctedErrorRate=0.105 corMinCoverage=0 genomeSize=330m”. The initial assembly had 763 contigs with N50 = 4.83 Mb (N50 number = 23).

### Hi-C data and scaffolding

Nuclei isolated as described [28] from both germline (testis) and somatic cells (5-day, 32-64 cell embryos) were used to generate Hi-C libraries using an adapted protocol [79,80]. The Hi-C data were used to further scaffold the initial contigs into chromosomes. Briefly, we first scaffolded the 763 contigs into 570 scaffolding (N50 = 10.24 Mb; N50 number = 13) using SALSA [81]. We then divided these scaffolds into 24 groups that correspond to the 24 germline chromosomes based on the contact information using Juicer and Juicebox [82]. For each chromosome group, we then mapped all original PacBio reads (360X) using MashMap [83] to retrieve all reads that can be uniquely mapped to the chromosome. The PacBio reads for each chromosome were then independently assembled using Canu, resulting in only 1 or 2 dominant contigs per chromosome for 20 chromosomes (the other 4 chromosomes have 3 or 4 large contigs). Each chromosome assembly was further evaluated manually using Juicer and Juicebox and scaffolded when necessary. The final assembly contains 24 scaffolds (N50 = 12.43 Mb; N50 number = 10) and 84 unplaced small contigs (2.3 % of the genome).

### Genes and their tissue-specific expression analysis

We mapped our previously defined transcripts [16] to define genes in the new genome assembly. Over 99.9% (61,766/61,820) of the genome-based transcripts and over 95.7% (59,965/62,645) *de novo* assembled transcripts can be mapped to the new genome, suggesting the current genome assembly is highly complete. To gauge the tissue-specific expression of the genes, we used RNA-seq data from 9 broadly defined tissues from *Ascaris*, including testis, ovary, zygote (zygote1-4), early embryos (24hr – 64hr), late embryos (96hr – 7day), larvae (L1 -L3), carcass, muscle and intestine [16,22,78]. The tissue-specific expression was scored by comparing each gene’s RNA level (rpkm) in the tissue where this gene is mostly highly expressed (has the highest rpkm) to the other 8 tissues. The expression of the gene in the tissue is considered as tissue specific (score = 100) if its expression is 4-fold higher than in any of the other 8 tissues. The fewer the number of tissues that meet the 4x expression cutoff the less tissue-specific expression the gene is. Genes with expression <4X higher than in all other tissues are considered as genes without tissue specific expression (score = 0); these genes are most likely house-keeping genes, particularly if their overall expression level is high.

### Repetitive sequence identification

Repetitive sequences were identified using a combination of homology-based and *de novo* approaches, including RepeatMasker [84], LTRharvest [85], RepeatScout [86], RepeatExplorer [87], and dnaPipeTE [88]. The repeats library was filtered for redundancy with CD-hit.

### Antibodies and immunohistochemistry

Monoclonal histone antibodies used (H3K9me2, H3K9me3, H3K36me3, H3K27me3, H3K4me3, and H4K20me1) were from Hiroshi Kimura [89,90] and have been previously validated for use in the modENCODE project for *C. elegans* and other organisms (see antibody validation database: http://compbio.med.harvard.edu/antibodies/). The *C. elegans* LGG-1 antibody was provided by Thorsten Hoppe [91] and detects both the non-lipidated (LGG-1-I) and the lipidated (LGG-1-II) forms of LGG-1. Commercial antibodies were obtained for rabbit Ubiquityl-Histone H2A (Lys119) (Cell Signaling D27C4), rabbit H3S10P (Millipore 06-570), and Nuclear Pore Complex Proteins (Abcam Mab414). *Ascaris* embryo immunohistochemistry was carried as described [28,78] using a modified freeze-crack method to permeabilize and fix embryos with some modifications. Briefly, decoated embryos were suspended in 50% methanol and 2% formaldehyde solution and were frozen and thawed 3 times using a dry ice/ethanol bath. The embryos were re-hydrated with 25% methanol in PBS pH 7.4 for 1 min. After washing twice with PBS pH 7.4, the embryos were suspended in blocking solution (0.5% BSA in PBS pH 7.4) for 1 hr at room temperature, followed by overnight incubation in primary antibodies at 4 °C, and then a 2 hr incubation in secondary antibodies (Invitrogen) at room temperature. Nuclei were stained with DAPI and slides mounted in anti-fade medium (Invitrogen).

### DNA FISH Probes

Telomere (5’-Quasar 570-TTAGGCTTAGGCTTAGGCTTAGGC) and 120 bp repeat (5’-Fam-CGAATAAATCCCAATTGCAG) oligonucleotide probes were obtained from LGC Biosearch (Petaluma, CA). For unique sequence FISH, 6-10 kb PCR products covering a total of 12-20 kb were generated for both retained and eliminated genome regions adjacent to internal and subtelomeric chromosome breaks. PCR primers were designed using the Oligo 7 software and PCR was performed using Long-Amp Taq DNA polymerase (New England Biolabs) following the manufactures directions. PCR products for a region were pooled, purified with SPRI beads (Beckman Coulter) and labeled with Gold 550 dUTP (Enzo) or Red 650 dUTP (Enzo) by nick translation using the Nick Translation DNA Labeling System 2.0 kit (Enzo). Labeled probes were then purified using Micro Bio-Spin® Columns with Bio-Gel® P-30 (BioRad).

### DNA Fish

A freeze-crack method was used to permeabilize [28,78] and fix the embryos in methanol/acetic acid (3:1). Methanol/acetic acid-fixed embryos were progressively rehydrated with 90%, 70%, 50% and 25% MeOH in PBS pH 7.4 for 10 minutes each at room temperature. Embryos were then washed once with PBS pH 7.4 and treated with 100 ug/mL RNase A in PBS pH 7.4 for 30 min at 37°C, followed by incubation in signal enhancer solution (Invitrogen) for 30 min at room temperature. Next, embryos were re-suspended in blocking solution (1% BSA in PBS pH 7.4) for 1 hr at room temperature and re-fixed in fresh 3.7% paraformaldehyde/1X PBS for 10 min at room temperature. Embryos were then washed with PBS pH 7.0, 2X SSC, and then incubated for 2 h at 37°C in pre-hybridization buffer (10% formamide in 2X SSC). Following pre-prehybridization, embryos were denatured in denaturation buffer (oligonucleotide probe buffer = 50% formamide in 2X SSC; Nick-translated probe buffer = 70% formamide in 2X SSC) for 5 min at 75°C. Oligonucleotide hybridization was carried out with 400 nM probes (telomere and 120 bp repeat) in hybridization buffer 1 (10% formamide/10% dextran sulfate/0.5 ug/uL salmon sperm DNA/0.2% SDS/2X SSC) overnight at 37°C. Duplex DNA probes were denatured for 5 min at 75°C, placed on ice for 2 minutes, pre-warmed at 37°C, and hybridization was carried out with 50 ng of nick-translated probe in hybridization buffer 2 (50% formamide/10% dextran sulfate/0.5 ug/uL salmon sperm DNA/0.2% SDS/2X SSC) overnight at 37°C. Post-hybridization washes for oligonucleotide probes were done in 2X SSC/0.1% Nonidet-P40 at 37°C for 5 min and for unique probes in 0.4X SSC at 45 °C for 15 min followed by 2X SSC/0.1% Nonidet-P40 at room temperature for 10 min. Nuclei were stained with DAPI, slides mounted in anti-fade medium (Invitrogen), and kept in the dark at room temperature for 24 hr before imaging.

### Image acquisition

*Ascaris* embryo immunohistochemistry and FISH images were acquired on an Applied Precision DeltaVision microscope, using a 60X immersion objective and FITC/Cy3/Cy5/DAPI excitation filter sets. Images were deconvolved with Applied Precision’s Softworx software and analyzed using Fiji software.

### Preparation of Ascaris embryos for electron microscopy

Embryos were prepared for electron microscopy using high pressure freezing and freeze substitution essentially as described [67] with the following modifications. Intact 4 cell and 8-16 cell embryos were collected as described above, mixed with 15% dextran, loaded into 100 um or 200 um aluminum planchettes, and then rapidly frozen under high pressure in a Wohlwend Compact 02 High Pressure Freezer (Technotrade International). The frozen samples were then freeze substituted in 2% OsO4 and 0.2% uranyl acetate in acetone for three days at −80°C, rinsed in acetone, warmed to −20°C, 4°C and then room temperature and embedded in Epon resin over 4 days. For immunolabeling, the frozen samples were freeze substituted in 0.1% glutaraldehyde and 0.01% uranyl acetate in acetone and embedded in Lowicryl HM20.

### Serial section electron microscopy, immuno-EM and electron tomography

Serial, 100-200 nm thick sections were cut using a Leica Ultracut UCT microtome. Sections were collected onto formvar-coated, copper slot grids and post stained with 2 % uranyl acetate and Reynolds lead citrate. The grids with serial sections were imaged using a Tecnai T12 microscope to identify embryos with mitotic cells, track the spindle through serial sections and to confirm the cell stage of a particular embryo.

For immuno-electron microscopy, serial, thin (80nm) sections were collected onto formvar-coated nickel slot grids. The grids were incubated in a blocking buffer containing 0.1% or 0.2% nonfat dry milk dissolved in PBS/Tween20, followed by incubation with the H3S10P antibody at 1:10 or 1:50 dilution. Control grids were incubated in blocking buffer without the primary antibody. The grids were rinsed and incubated with a rabbit secondary antibody conjugated to 15 nm gold at a 1:20 dilution in blocking buffer (Ted Pella, Redding, CA). Grids were post stained with 2% uranyl acetate and Reynolds lead citrate and imaged in a Tecnai T12 microscope. Images from serial sections were collected to confirm labeling on specific organelles.

Tomography was performed as essentially as described [68]. Briefly, grids containing serial, 200nm thick sections were imaged using a Tecnai F30 (Thermo Fisher Scientific, Waltham, MA) operating at 300kV. Montaged images were collected using a Gatan OneView camera (Gatan, Inc. Pleasanton, CA) to record images at a 1.7nm pixel size over an area approximately 7um x 7um. Single axis tilt series were collected over a +60° to −60° range at 1° increments using the SerialEM image acquisition program [92]. The tilted views were aligned using local patch tracking and tomograms generated using the IMOD 4.9 software package (https://bio3d.colorado.edu/imod/) [93,94].

Tomograms were displayed and microtubules modeled using the 3dmod program of the IMOD software package. The image slicer window was used to orient a slice of image data to follow and track individual MTs along their lengths. The model contours were projected in 3D and rotated to study the spindle MT organization during DNA elimination mitosis.

## Acknowledgements

We thank Bruce Bamber, Jeff Myers, and Routh Packing Co. for their support and hospitality in collecting *Ascaris* material. We thank Garry Morgan and Thomas Giddings for assistance with specimen preparation and immuno-EM experiments, the University of Colorado at Boulder EM Services Core Facility, and Thorsten Hoppe for generously providing the *C. elegans* LGG-1 antibody. We thank Dick Mcintosh for his interest, insight, and feedback on this work, Peter Carlton for bringing to our attention the issue of independent sex chromosomes in somatic cells of *Parascaris*, and Rachel Patton McCord for interpretation of the Hi-C data. This work was supported by NIH grants to R.E.D. (AI114054) and J.W. (AI125869).

## Author contributions

JW and RED designed the project; JW and RED prepared agarose plugs and megabase size genomic DNA; YK and JW carried out the Hi-C experiments; JW and MZ carried out bioinformatic analyses; GMBV designed and carried out the DNA FISH and IHC; EOT designed and carried out the EM analyses; JW, GMBV, MZ, EOT, and RED analyzed data; JW, EOT, and RED wrote the manuscript.

## Declaration of Interests

The authors declare no competing interests.

---

**Supplementary Figure S1.**
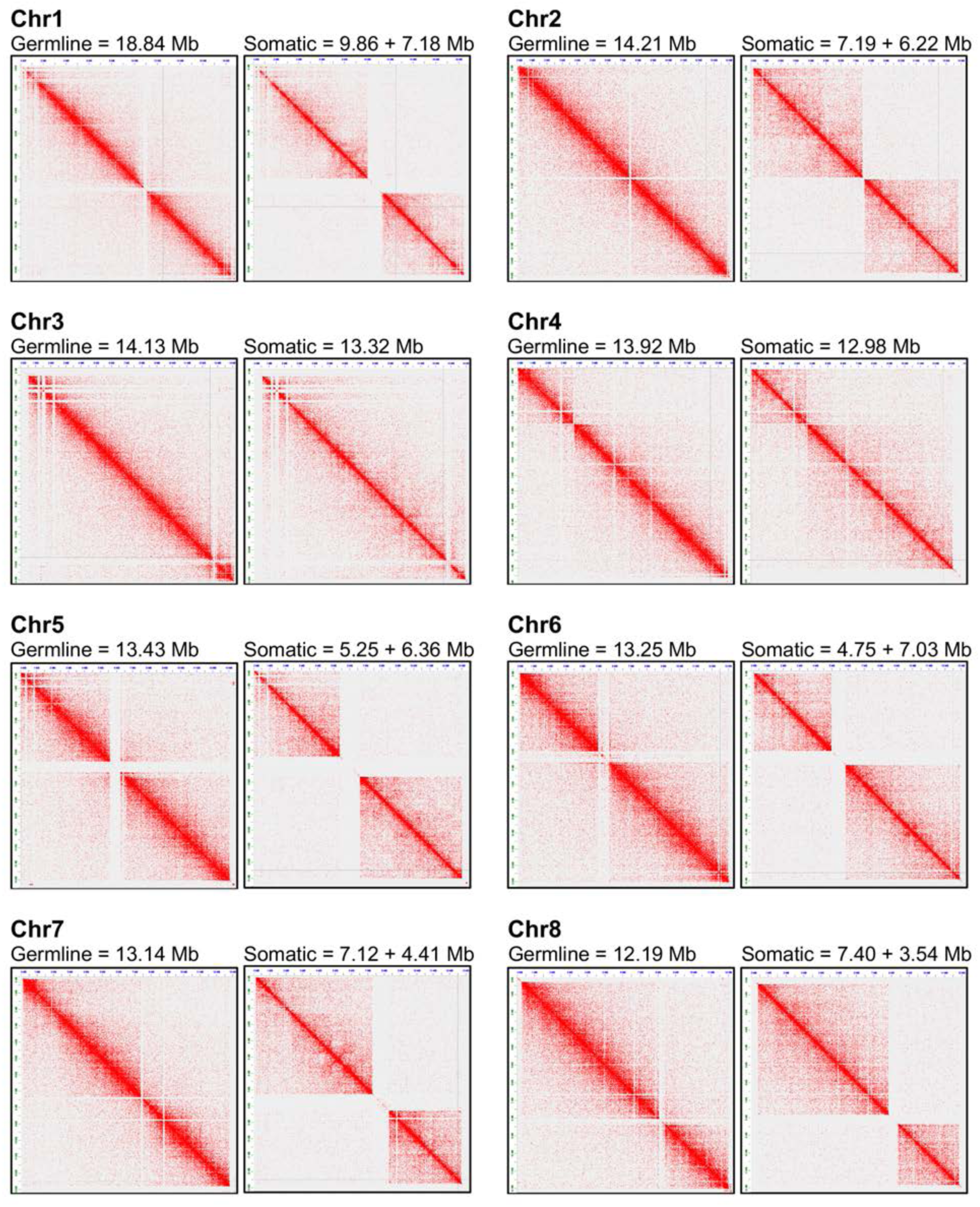
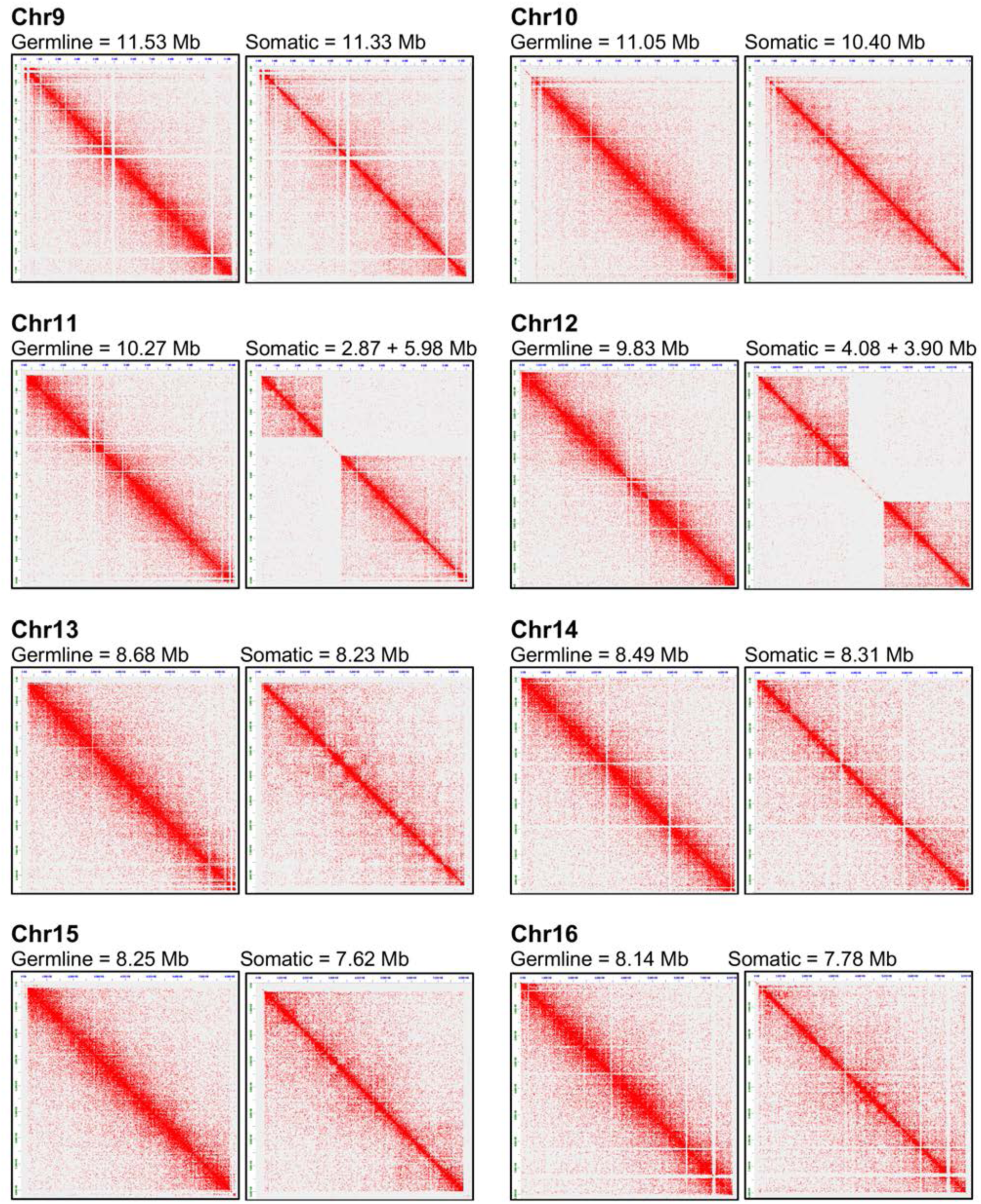
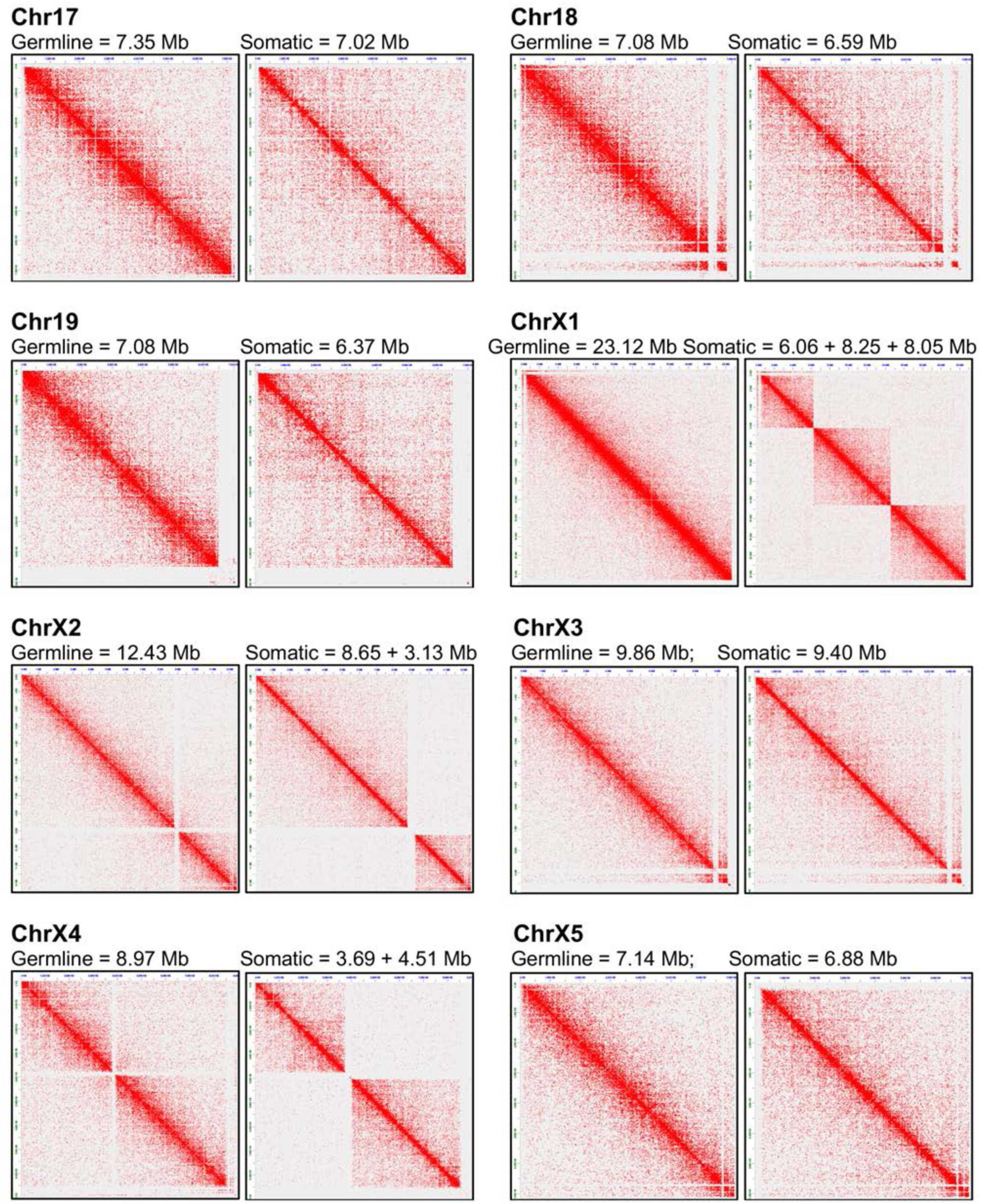
Hi-C interaction changes between germline and somatic cells in all 24 *Ascaris* germline chromosomes. Germline Hi-C data is derived from testis and somatic data is from 32-64 cell (post-DNA elimination, 5 day) embryos. This figure is divided into three pages.

**Supplementary Figure S2.**
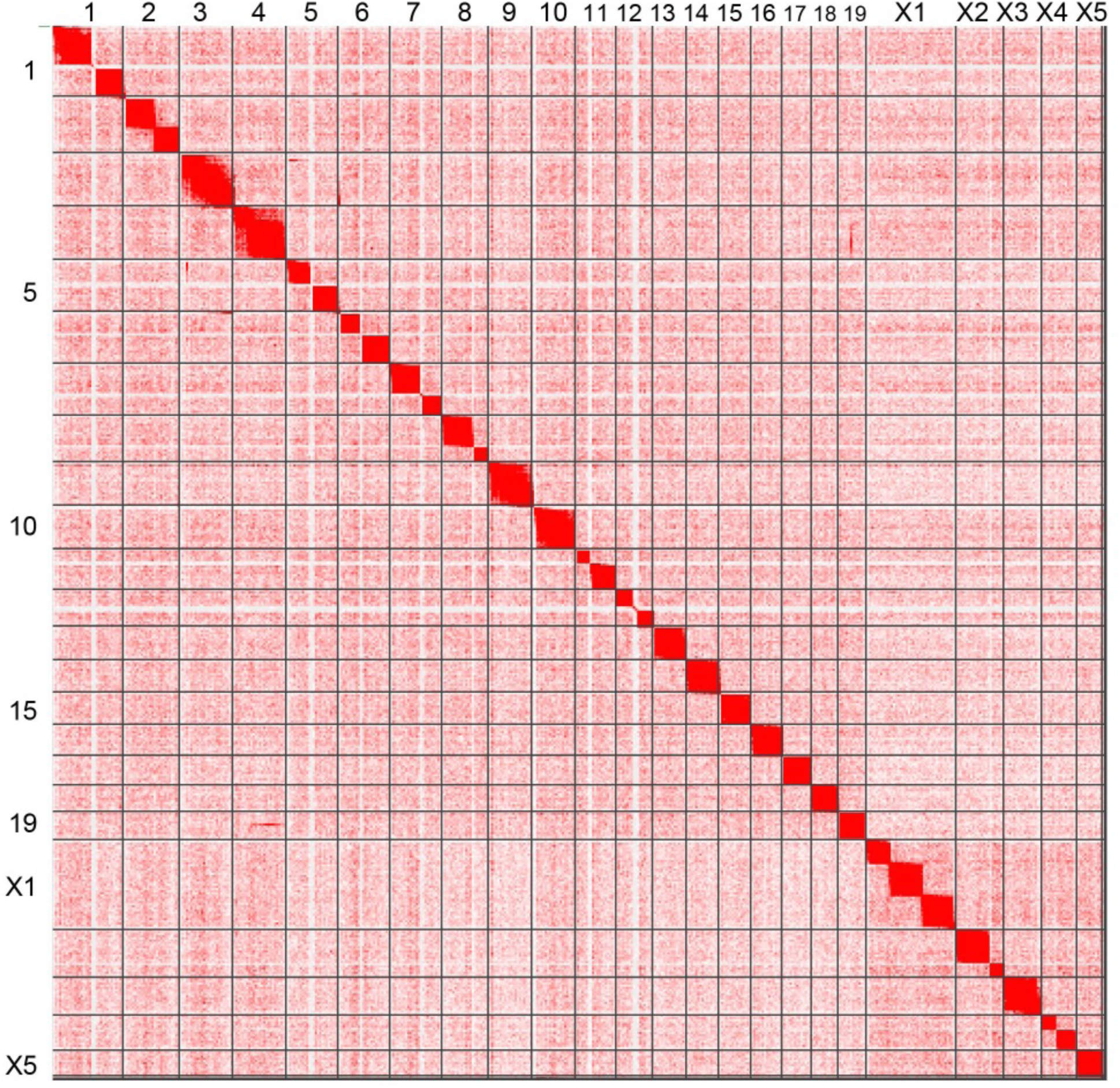
Hi-C data showing the interactions between *Ascaris* autosomes (1-19) and sex chromosomes (X1-X5) in 5 day (32-64 cell) embryos.

**Supplementary Figure S3.**
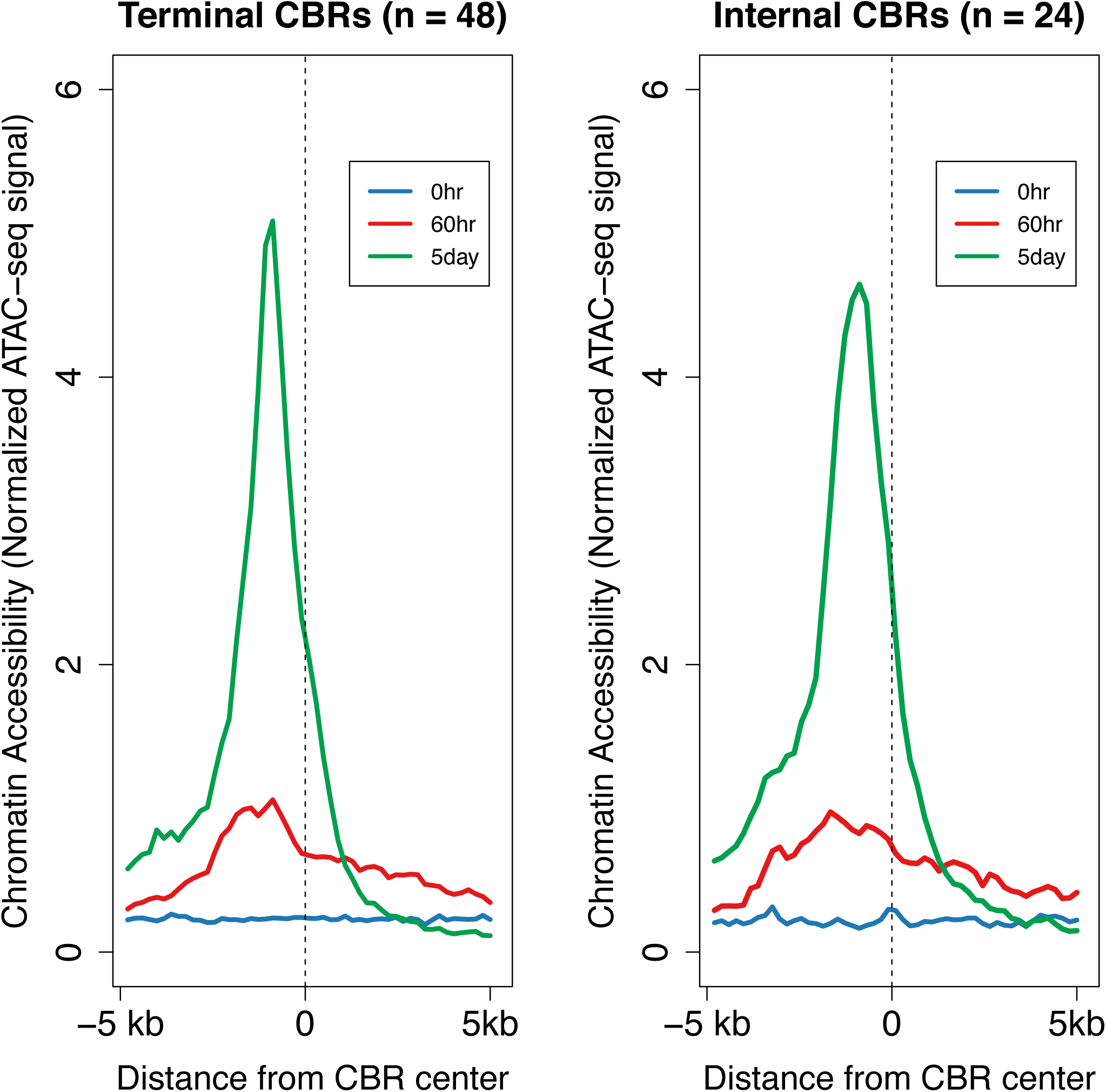
Increased chromatin accessibility is associated with both terminal and internal CBRs for *Ascaris* DNA elimination. Illustrated are CBRs with 5-kb flanking regions. Upstream of the center of the CBRs (−5 kb to 0) is retained DNA, while downstream (0 to 5 kb) is eliminated DNA. DNA accessibility increases from 0 hr embryos (1-cell stage) to 60 hr embryos (4-cell; immediately prior to DNA elimination) and increases and persists at 5 day embryos (32-64 cell). Note that the open regions correspond exactly to the CBRs defined by the regions where DNA breaks and telomere addition occurs.

**Supplementary Figure S4 (related to Figure 3).**
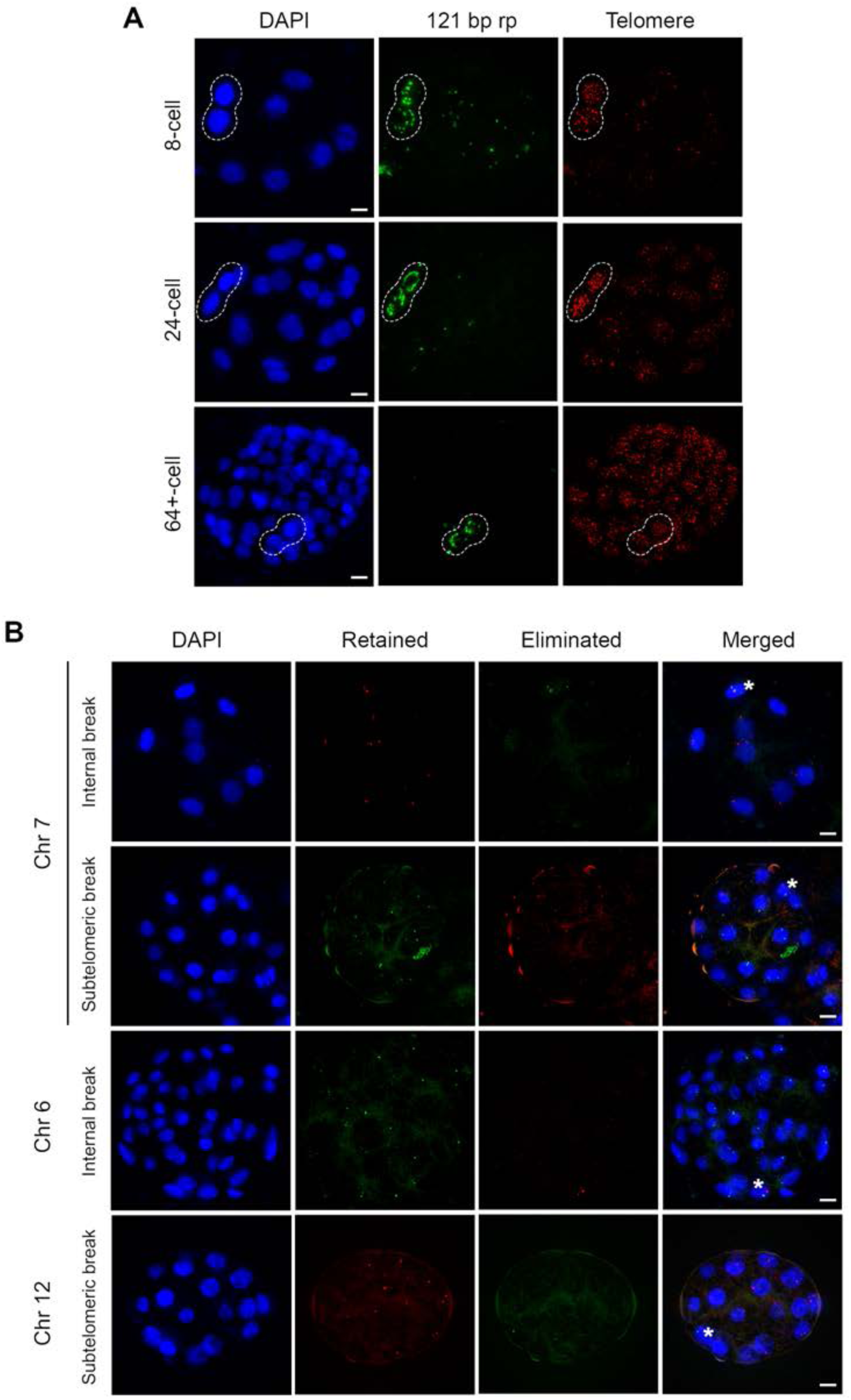
DNA FISH illustrates retention and loss of DNA following DNA elimination. **(A)** Single-channel images of DNA FISH for telomeric and 120 bp repeats shown in Figure 3 C-E. Dotted circle indicate germline cells. **(B)** DNA FISH for eliminated and retained regions of internal and subtelomeric chromosome breaks in 8- to 32+-cell embryos. Note probes corresponding to eliminated sequences are only detected in germline cells (*) while probes for retained sequences show signal in somatic nuclei as well. Scale bars = 5 µm.

**Supplementary Figure S5 (related to Figure 4).**
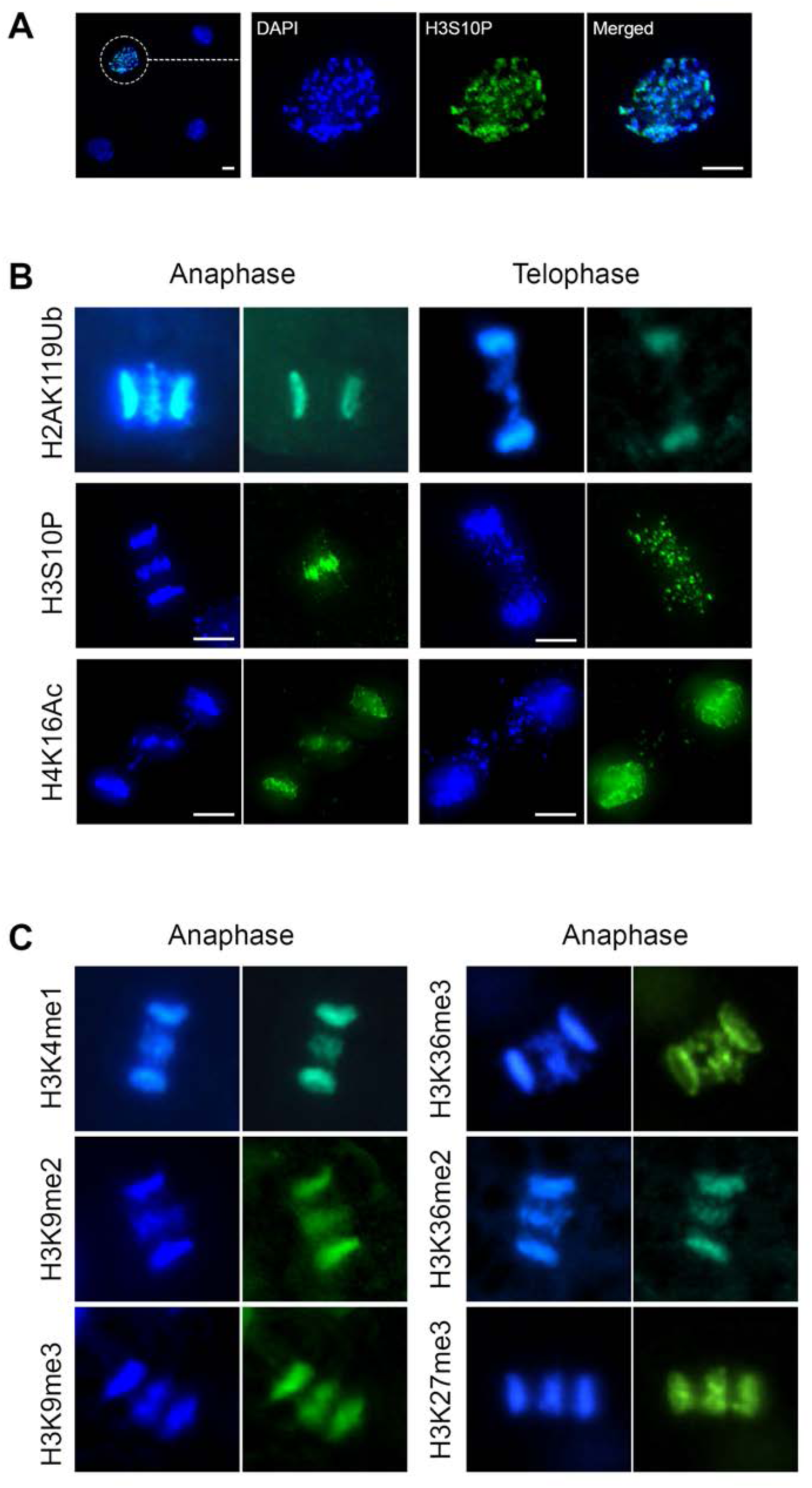
Histone staining of *Ascaris* prophase and DNA elimination anaphase and telophase chromosomes. **(A)** H3S10P staining on prophase chromosomes is not uniform. **(B)** H2AK119Ub is reduced in DNA that is going to be eliminated at DNA elimination telophase. H3S10P staining of eliminated DNA seen in anaphase persists in telophase while H4K16ac, like H3K4me3 (see Fig. 4), is removed at telophase during DNA elimination, when the DNA is incorporated into vesicles. (**C)** Other histone modification marks equally stain retained and eliminated DNA at DNA elimination anaphase. Scale bars = 5 µm.

**Supplementary Figure S6 (related to Figure 6).**
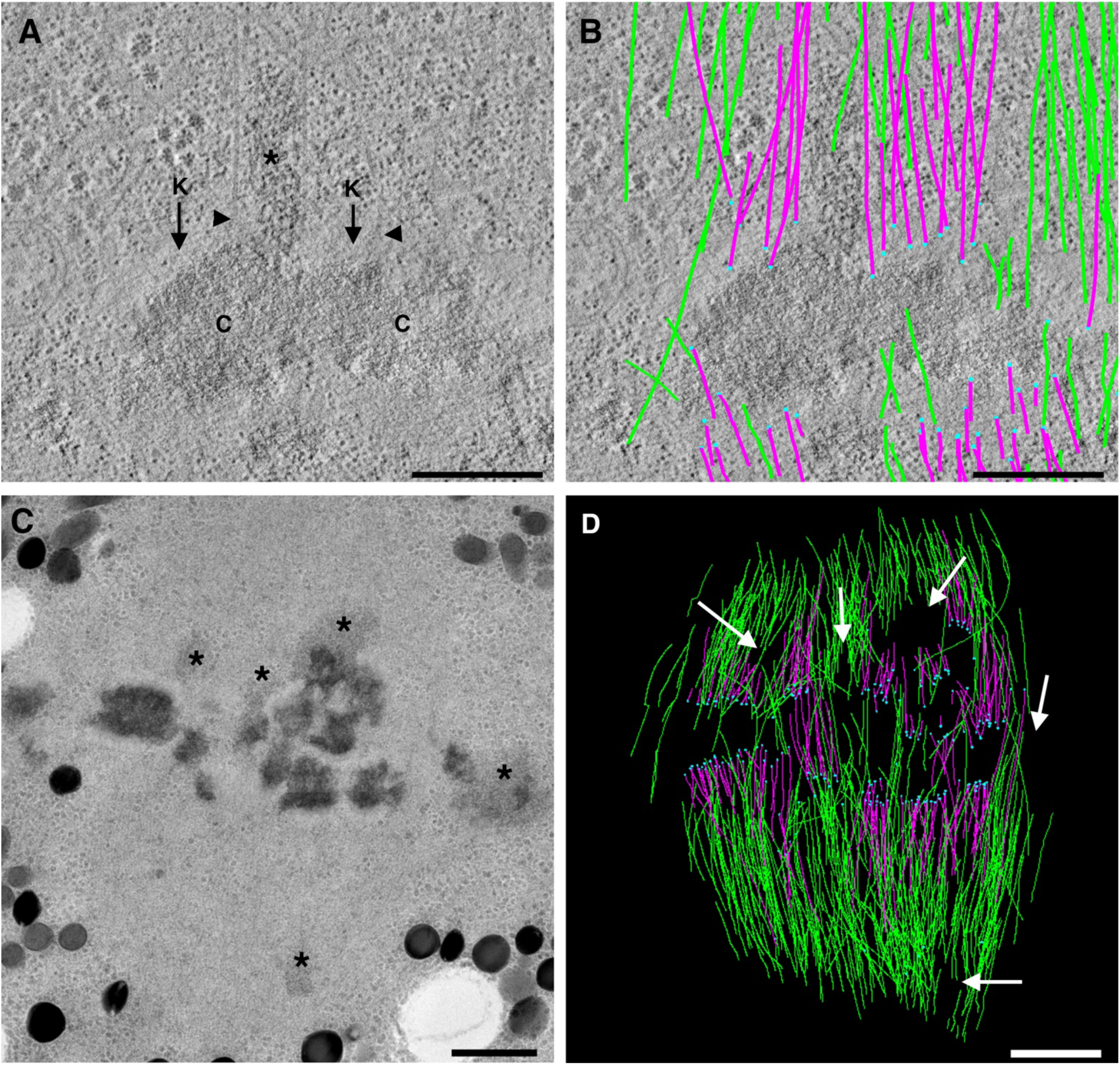
Ultrastructural tomographic analysis of a DNA elimination metaphase. Note the retained DNA is bound by kinetochores and kMTS while DNA that will be eliminated is not. **(AB)** Late prometaphase of a DNA elimination mitosis at the 4-cell stage. Only condensed chromosomes assemble kinetochores and have KMTs. **(A**) Tomographic slice showing two condensed chromosomes (C) with a mass of diffuse chromatin adjacent to them (*). The condensed chromosomes assemble kinetochores (k) and have KMTs (arrowheads). Serial, tomographic slices showing modeled MTs in this region are shown in Movie S3. Bar= 500nm. **(B)** 3D model shows numerous KMTs (pink) making end-on attachments (blue spheres) to the kinetochores of each chromosome but not at the diffuse chromatin. Other nonKMTs traverse the region (green). These nonKMTs either go out of the volume reconstructed. Bar= 500nm. **(C-D)** Another region of the late prometaphase spindle showing KMTs at chromosomes but not at the diffuse chromatin (see also Figure 6D). **(C)** Overview of one of three 200nm thick sections imaged for tomography shows darkly-stained, condensed chromosomes at the metaphase plate and lighter stained chromatin masses in the spindle proper and near the metaphase plate (*’s). Bar = 1um. **(D)** Projected 3D model built from 3 serial sections show KMTs (pink) with ends marked with blue spheres. NonKMTs are shown in green. White arrows point to regions where diffuse chromatin lays; MTs surround but do not make end-on contact with it (see also Movie S4). Bar = 1um.

**Supplementary Figure S7.**
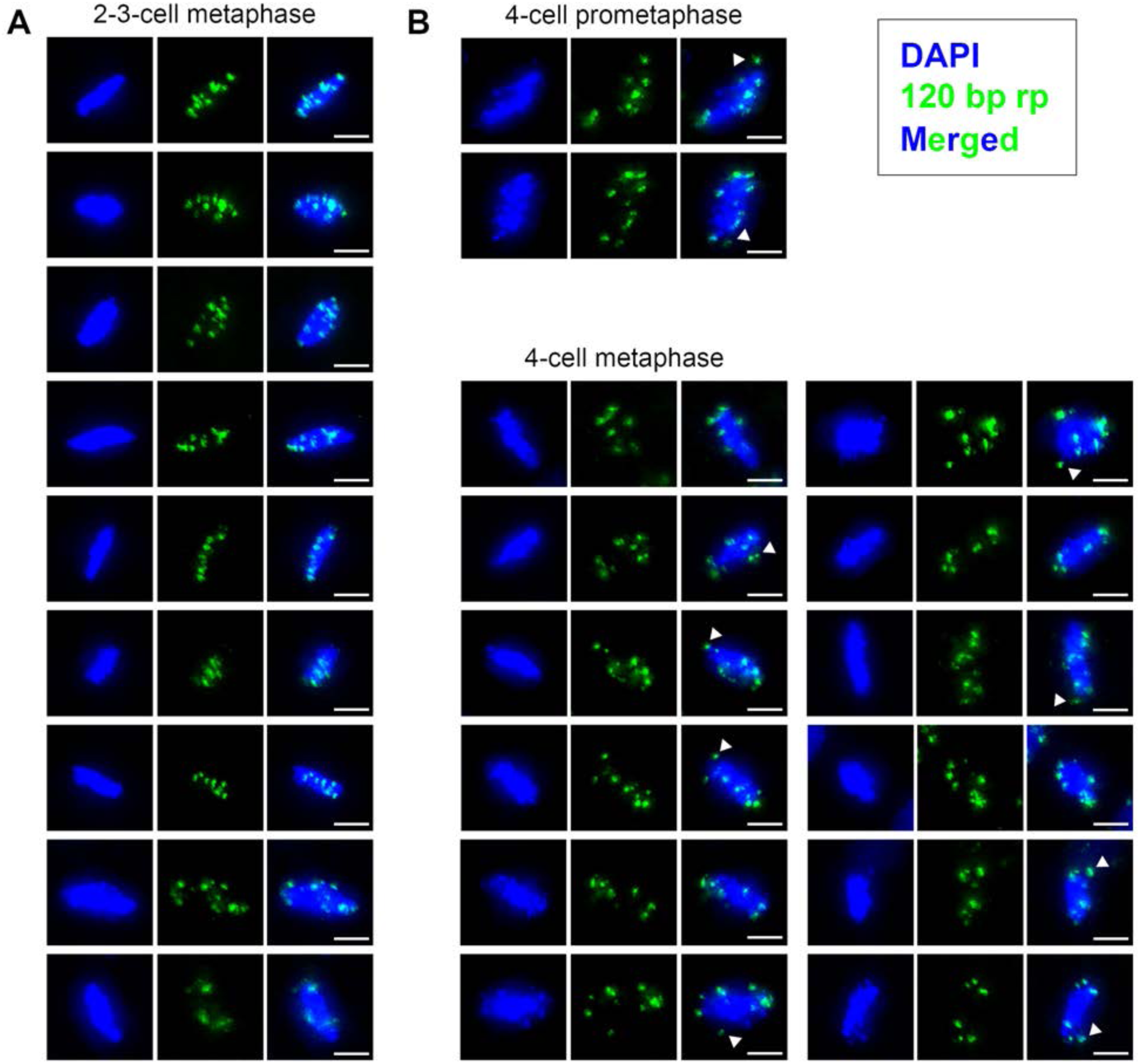
DNA FISH for the 120 bp repeat during mitotic metaphases shows the repeat at the boundary of the metaphase plate during a DNA elimination. **(A-B)** Differential distribution of 120 bp repeat in regular versus DNA elimination mitosis. **(A)** Metaphase of a normal mitosis (non-DNA elimination) at 2-3 cell stage. The 120 bp repeat is coincident with DNA and located all through the metaphase plate. **(B)** At prometaphase and metaphase of 4-cell embryo DNA elimination mitosis, the 120 bp repeat locates mainly to the boundary of the metaphase plate and shows delayed congression (arrowheads). Scale bars = 5 µm.

## Supplementary Movie Legends

Movies available at https://drive.google.com/open?id=1RN1VErbG9iSBG4ahJ-qzgeVgsly7EWKj

**Movie S1. Serial, tomographic slices and a projected 3D model from part of the spindle of an *Ascaris* DNA elimination mitosis in prometaphase** (the cell shown in Figure 5D). The tomographic volume was built from two, serial 200nm thick sections comprising a volume ~7um x 7um x 400nm. KMTs are purple (n=115), blue spheres mark KMT ends at the chromosome kinetochores, Mts that do not end on chromosomes or go out of the volume of the reconstruction are green (n=345). Bar = 1µm.

**Movie S2. Serial, tomographic slices of a partial volume from the spindle shown in movie 1.** Model contours track the trajectories of MTs surrounding diffuse chromatin in the spindle proper (as marked by a white arrow in Figure 6-A and B). MTs do not end on this kind of chromatin but make close, lateral contacts. Bar = 50 nm

**Movie S3. Serial, tomographic slices of a partial volume from the spindle shown in Figure 6A.** Two condensed chromosomes have kinetochores and KMTs (purple). Adjacent to these chromosomes is an object containing diffuse chromatin that does not bind KMTs. Numerous nonKMTs can be seen throughout the volume (green) Bar = 250 nm

**Movie S4. Serial, tomographic slices and projected 3D model from a different region of the spindle from an *Ascaris* DNA elimination mitosis in prometaphase.** The tomographic volume was built from three, serial 200nm thick sections comprising a volume ~7um x 7um x 400nm. KMTs are purple (n=187), blue spheres mark KMT ends at the chromosome kinetochores, MTs that do not end on chromosomes or go out of the volume of the reconstruction are green (n=476). These MTs do not end on diffuse chromatin but have close, lateral contact with it. Bar = 1µm.

## Notes

https://drive.google.com/open?id=1RN1VErbG9iSBG4ahJ-qzgeVgsly7EWKj

### Competing Interest Statement

The authors have declared no competing interest.

